# A possible coding for experience: ripple-like events and synaptic diversity

**DOI:** 10.1101/2019.12.30.891259

**Authors:** J Ishikawa, T Tomokage, D Mitsushima

## Abstract

The hippocampal CA1 is necessary to maintain experienced episodic memory in many species, including humans. To monitor the temporal dynamics of processing, we recorded multiple-unit firings of CA1 neurons in male rats experiencing one of four episodes for 10 min: restraint stress, social interaction with a female or male, or observation of a novel object. Before an experience, the neurons mostly exhibited sporadic firings with some synchronized (≈ 50 ms) ripple-like firing events in habituated home cage. After experience onset, restraint or social interaction with other rats induced spontaneous high-frequency firings (super bursts) intermittently, while object observation induced the events inconsistently. Minutes after experience initiation, CA1 neurons frequently exhibited ripple-like firings with less-firing silent periods. The number of ripple-like events depended on the episode experienced and correlated with the total duration of super bursts. Experience clearly diversified multiple features of individual ripple-like events in an episode-specific manner, sustained for more than 30 min in the home cage.

*Ex vivo* patch clamp analysis further revealed experience-promoted synaptic plasticity. Compared with unexposed controls, animals experiencing the female, male, or restraint episodes showed cell-dependently increased AMPA- or GABA_A_ receptor– mediated postsynaptic currents, whereas contact with a novel object increased only GABAergic currents. Multivariate ANOVA in multi-dimensional virtual space revealed experience-specific super bursts with subsequent ripple-like events and synaptic plasticity, leading us to hypothesize that these factors are responsible for creating experience-specific memory. It is possible to decipher encrypted experience through the deep learning of the orchestrated ripple-like firings and synaptic plasticity in multiple CA1 neurons.

## Introduction

The hippocampus is a primary site for episode-like memory development (Scoville & Milner, 1957), known to process spatio-temporal information (Mitsushima et al, 2009; Wills et al, 2010) within a specific episode (Gelbard-Sagiv et al, 2008). The dorsal CA1 neurons may encode place and context when animals are exploring novel context (Tanaka et al, 2018), and temporal inactivation of firings before exploration of novel objects can impair test performance in a what-where-when episodic-like memory task (Drieskens et al, 2017). These observations indicate the importance of firing activity during or immediately after episodic experience, but specific firing patterns during the early learning period are not clear.

Emotions such as happiness, fear, and sadness influence the strength of a memory (Christianson et al, 1992; McGaugh et al, 2000; Richter-Levin & Akirav, 2003; LeDoux, 2000). Emotional arousal enhances learning via noradrenergic stimulation of the dorsal CA1 neurons to drive GluA1-containing AMPA receptors into the synapses (Hu et al, 2007). Tyrosine hydroxylase–expressing neurons in the locus coeruleus may mediate post-encoding memory enhancement with co-release of dopamine in the hippocampus (Takeuchi et al, 2016). However, conclusive evidence regarding whether emotional arousal affects CA1 neuron firing and the associated temporal dynamics is lacking in freely behaving animals.

The first aim of this study was to examine the temporal dynamics of CA1 neural activity in the early learning period in rats. The second aim was to examine differences in these firing patterns among different experiences. For this purpose, we used four emotionally distinct episodes and compared the associated early-learning processes. Temporo-spatial firing patterns by multiple neurons may orchestrate a possible code (Grinvald et al, 2003; van Hemmen & Sejnowski, 2006), so we also monitored changes in multiple-unit firings after various episodic experiences. Finally, by analyzing postsynaptic currents induced by a single vesicle of glutamate or GABA *ex vivo* (Sakimoto et al, 2019), we examined experience-specific plastic changes at excitatory/inhibitory synapses onto the CA1 pyramidal neurons.

## Materials and Methods

### Animals

Sprague–Dawley male rats (CLEA Japan Inc., Tokyo, Japan) were housed at 22°C with a 12-h light–dark cycle (lights on from 8:00 A.M to 8:00 P.M.). Rats were allowed at least 2 weeks of *ad libitum* food (MF, Oriental Yeast Co. Ltd., Tokyo, Japan) before surgery. Rats at age 15 to 25 weeks were used, and all experiments were conducted during the light cycle. These studies were reviewed and approved by the Yamaguchi University Graduate School of Medicine Committee of Ethics on Animal Experiments. All manipulations and protocols were performed according to the Guidelines for Animal Experiments at Yamaguchi University Graduate School of Medicine and in accordance with Japanese Federal Law (no. 105), Notification (no. 6) of the Japanese Government, and the National Institutes of Health Guide for the Care and Use of Laboratory Animals (NIH publications no. 85-23), revised in 1996.

### Surgery

Animals were anesthetized with sodium pentobarbital (50 mg/kg, intraperitoneal) and placed in a stereotaxic apparatus. Movable recording electrodes (Unique Medical Co., LTD, Japan) were chronically implanted above the hippocampal CA1 (posterior, 3.0– 3.6 mm; lateral, 1.4–2.6 mm; ventral, 2.0–2.2 mm) and fixed with dental cement. Rats were housed individually and excluded for analysis if cannulas or electrodes did not target the region.

### In vivo recording of multiple-unit firing activity

Neural signals were passed through a head amplifier and then into the main amplifier (MEG-2100 or MEG-6116; Nihon Kohden, Tokyo, Japan) through a shielded cable. Signals were band-pass filtered at 150 – 10 kHz and digitized using a CED 1401 interface controlled by Spike2 software (Cambridge Electronics Design, Cambridge, UK). Signal data were mostly sampled at 25 kHz, but a few were sampled at 17 kHz.

Isolation of single units was initially performed using the template-matching function in Spike2 software. As we reported previously (Ishikawa et al, 2015), all spikes used in subsequent analysis were clearly identified, with a signal-to-noise ratio of at least 3 to 1. Following the initial separation of spikes, we applied principal component analysis of the detected waveforms. Single units had to show cluster separation after their first three principal components were plotted (Fig. 1E). However, in this experiment, the sorting was not always reliable, especially in super bursts and ripple-like events, because one electrode recorded many units. Therefore, we analyzed all recording data as multiple unit activity.

**Figure 1.**
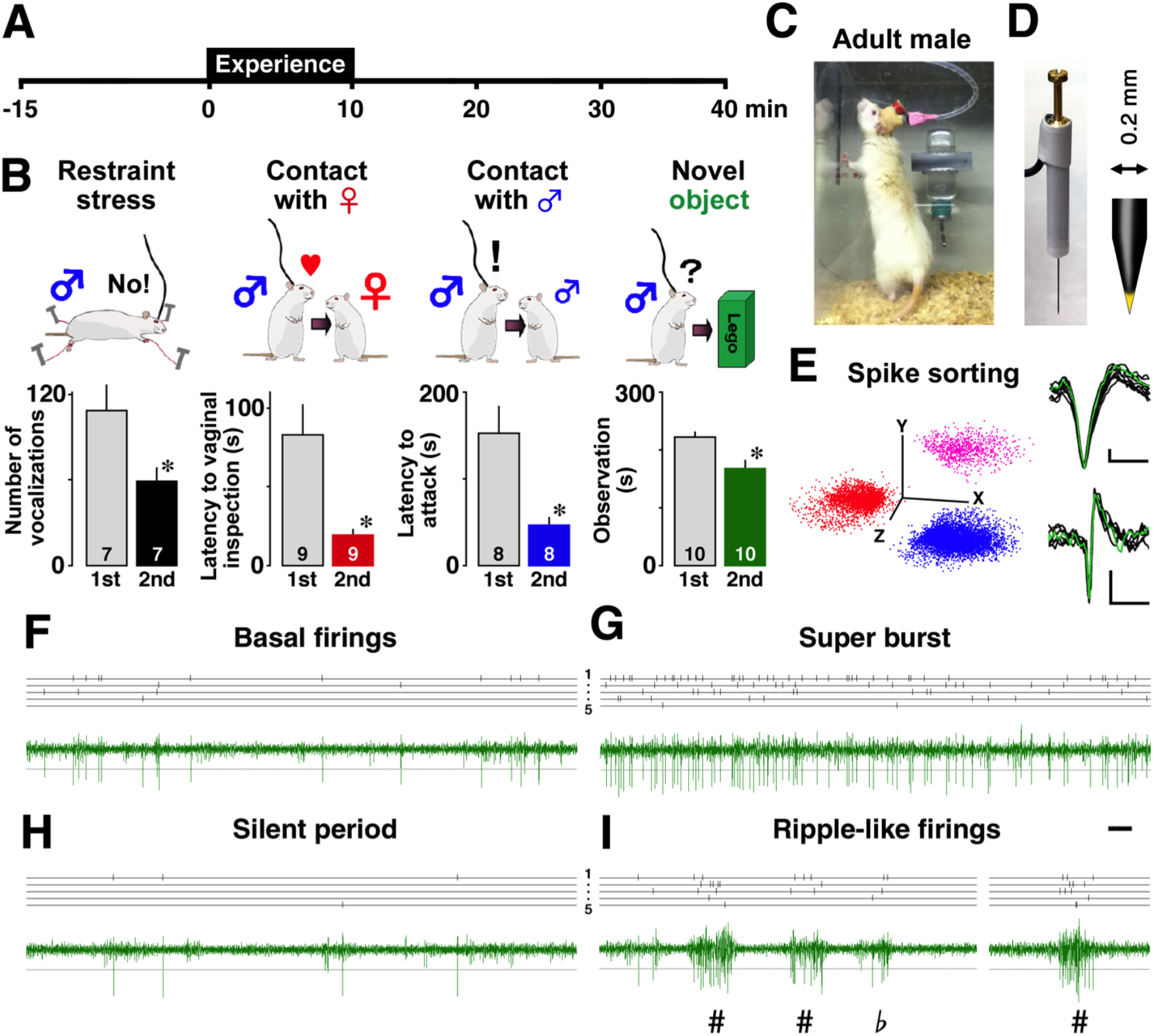
Experiences and extracted spike patterns. (A) Schedule of multiple-unit recording and 10-min experience. (B) To evaluate the experienced memory, behaviors during the first experience were compared with those during the second experience. (C) Multiple-unit spike activity was recorded in the CA1 neurons of adult male rats in their habituated home cage. (D) A picture of the recording electrode and enlarged view. (E) Examples of sorted spikes. Vertical bar = 0.2 mV; horizontal bar = 1 ms. Multiple features of individual spikes were plotted, and principal components were analyzed using Spike2 software. Examples of basal firings (F), super burst (G), silent period (H), and three ripple-like events (I) recorded from the same electrode (300–10 kHz). Upper bars indicate sorted spikes. # = ripple-like events; b = a sub-threshold event; horizontal bar = 50 ms. The number of rats in each group is shown at the bottom of each bar. \**P* < 0.05, \*\**P* < 0.01 *vs*. 1st experience. Error bars indicate ± SEM.

We identified a spontaneous event of super burst (Figs. 1G and 2) as one showing a higher firing rate for each point greater than 3 standard deviations (SDs) of the mean firing rate before the episodic experience. Figure 2F shows profile plots of individual super bursts over the threshold. A silent period (Figs. 1H and 3A) was one that showed a longer duration of inter-spike interval than 3 SD above the mean of the baseline. A ripple-like event was defined as one that involved long-lasting, high-frequency firing (> 10 ms) with a signal-to-noise ratio of at least 6 to 1 (Figs. 1I and 3A). Behavioral states during the ripple-like events were immobile eye-closed 16.0 ± 7.0 %, immobile eye-opened 59.7 ± 11.0%, or eye-opened moving 24.3 ± 11.3% (1631 events in 9 rats were analyzed).

**Figure 2.**
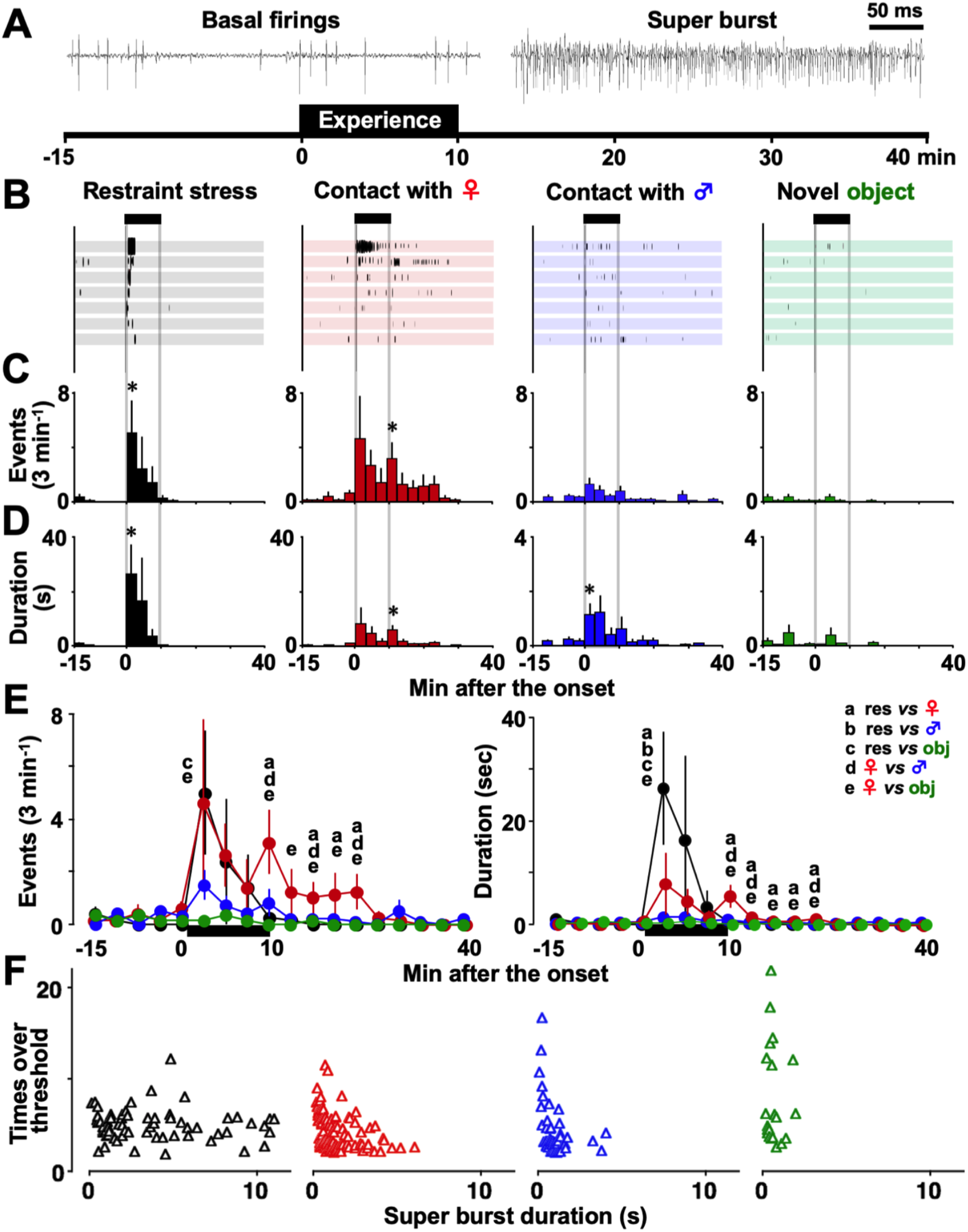
Super bursts and the experience-specific features. (A) An example of basal firings and experience-induced super burst of CA1 neurons. (B) Timeline plots of super bursts before, during, and after the experiences in individual rats. Restraint stress and contact with a female rat clearly induced many burst events. Contact with a male rat also induced some burst events, but the novel object inconsistently did so. (C) The incidence and total duration of the events (D) before, during, and after the experiences. (E) The incidence and total duration of super bursts was statistically different among groups. (F) Two-dimensional plots of individual super bursts during and after the experiences. A triangle indicates a single super burst event, showing significant experience-specificity (*P* < 0.001 in MANOVA). \**P* < 0.05, \*\**P* < 0.01 *vs*. before. Letters indicate *P* < 0.05 for the following comparisons: a, restraint *vs*. female; b, restraint *vs*. male; c, restraint *vs*. object; d, female *vs*. male; e, female *vs*. object. Error bars indicate ± SEM.

**Figure 3.**
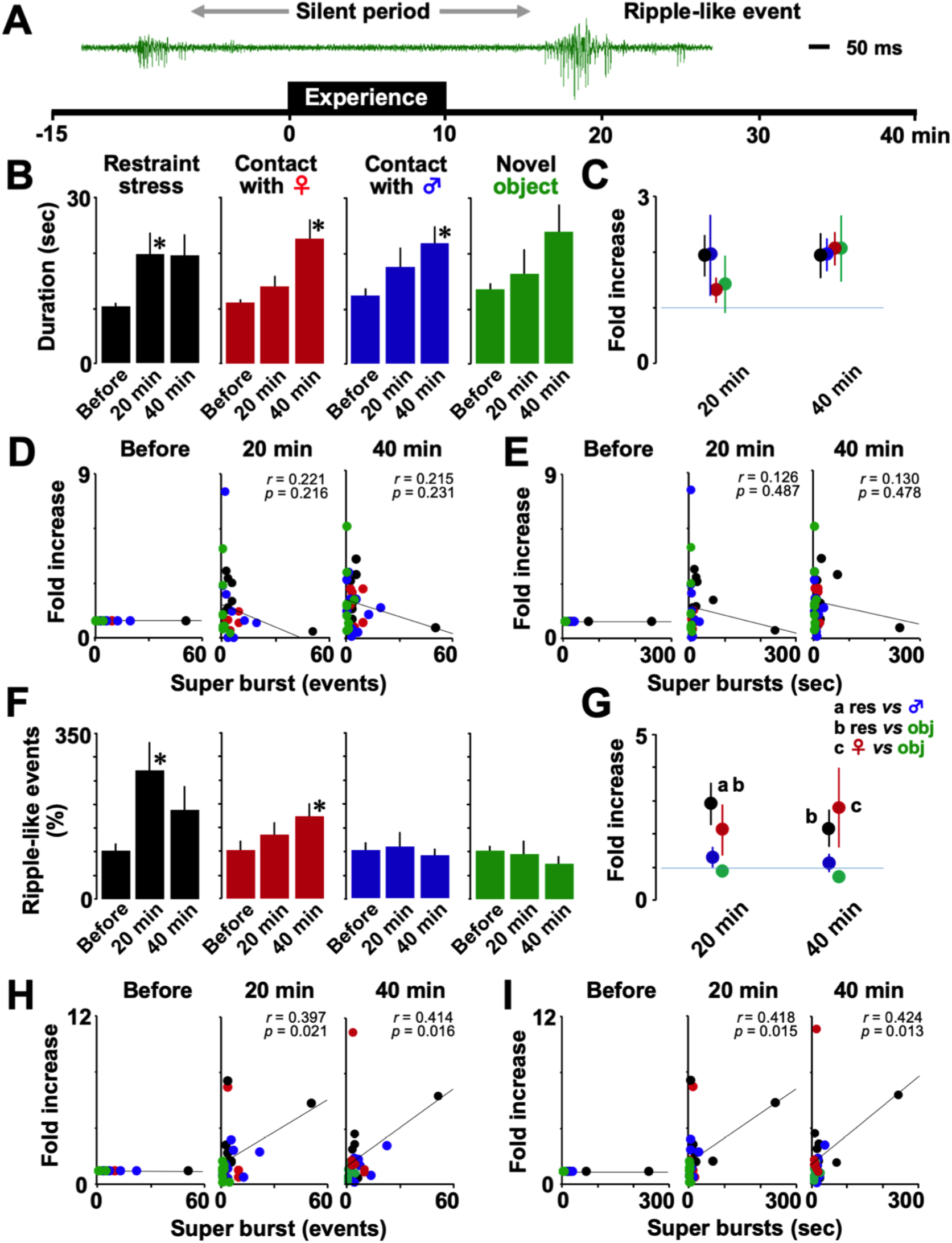
Silent periods and ripple-like events. (A) Representative examples of a silent period with a ripple-like event. (B) Restraint stress or contact with another rat significantly increased the duration of silent periods. (C) No significant difference was observed among the experiences. The duration was not correlated with the incidence (D) or total duration of super bursts (E). (F) Relative incidence of ripple-like events. Restraint stress or contact with a female rat increased the incidence of ripple-like events, while contact with a male or novel object did not affect the incidence. (G) Changes in the incidence of ripple-like events was experience-dependent. The number of ripple-like events was significantly correlated with the incidence (H) or the total duration of super bursts (I). \**P* < 0.05 *vs*. before. Letters indicate *P* < 0.05 for the following comparisons: a, restraint *vs*. male; b, restraint *vs*. object; c female *vs*. object. Error bars indicate ± SEM.

To detect ripple-like events, recorded signal (150–10 kHz) were filtered at 150– 300 Hz and 300–10 kHz. Using the 150–300 Hz signal, we calculated the root mean square (RMS) and the threshold for the event detection was set to +6 SD above the mean of the baseline. Moreover, the duration of a ripple-like event was defined from the time at the RMS reached +3 SD of baseline to the time at the RMS came down to the level. The calculated duration of the event (Fig. 4D) was shorter than the visual feature.

**Figure 4.**
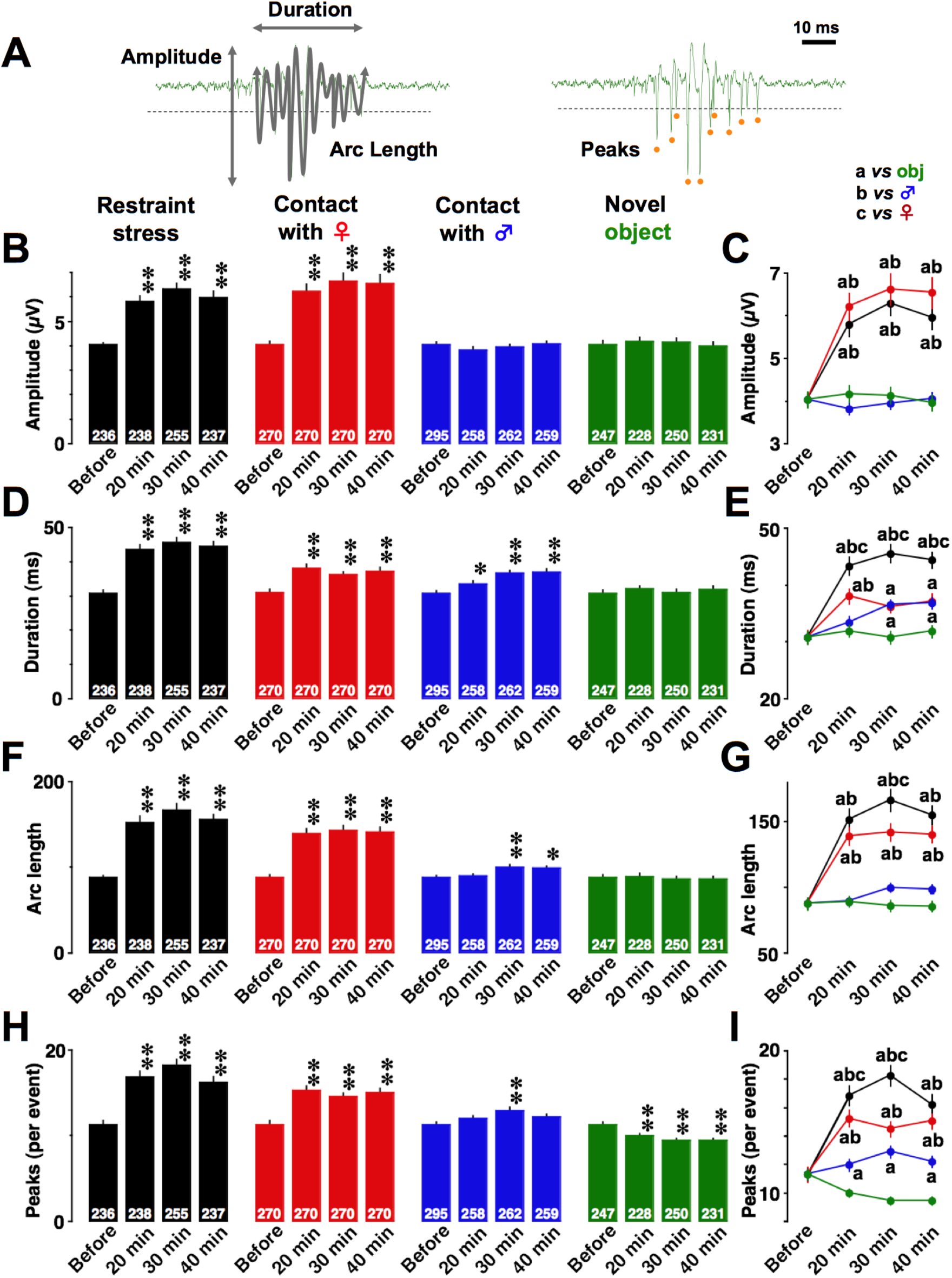
Four features of individual ripple-like events. (A) An example of ripple-like event with the amplitude, duration, arc length, and number of peaks. (B) Temporal changes in the amplitude of ripple-like events. (C) The amplitude comparisons among four experienced groups. (D) Temporal changes in the duration of ripple-like events. (E) The duration comparisons among four experienced groups. (F) Temporal changes in the arc length of ripple-like events. (G) The arc length comparisons among four experienced groups. (H) Temporal changes in the peaks of ripple-like events. (I) The peaks comparisons among four experienced groups. The number of cells in each group is shown at the bottom of each bar. \**P* < 0.05, \*\**P* < 0.01 *vs*. before. Letters indicate *P* < 0.05 for the following comparisons: a, *vs*. object; b, *vs*. male; c, *vs*. female. Error bars indicate ± SEM.

We used the 300–10 kHz signal for Figures 1 to 5. Although the ripple-like events (300–10 kHz) were mostly accompanied with the sharp-wave ripples (SWRs: 150–300 Hz), the shape of ripple-like events are much different from the SWRs. If a ripple-like event was not accompanied with SWR, we excluded the event for analysis. We used the 300–10 kHz signal to analyze amplitude, arc length, or peaks (Figs 4B, 4F, and 4H).

**Figure 5.**
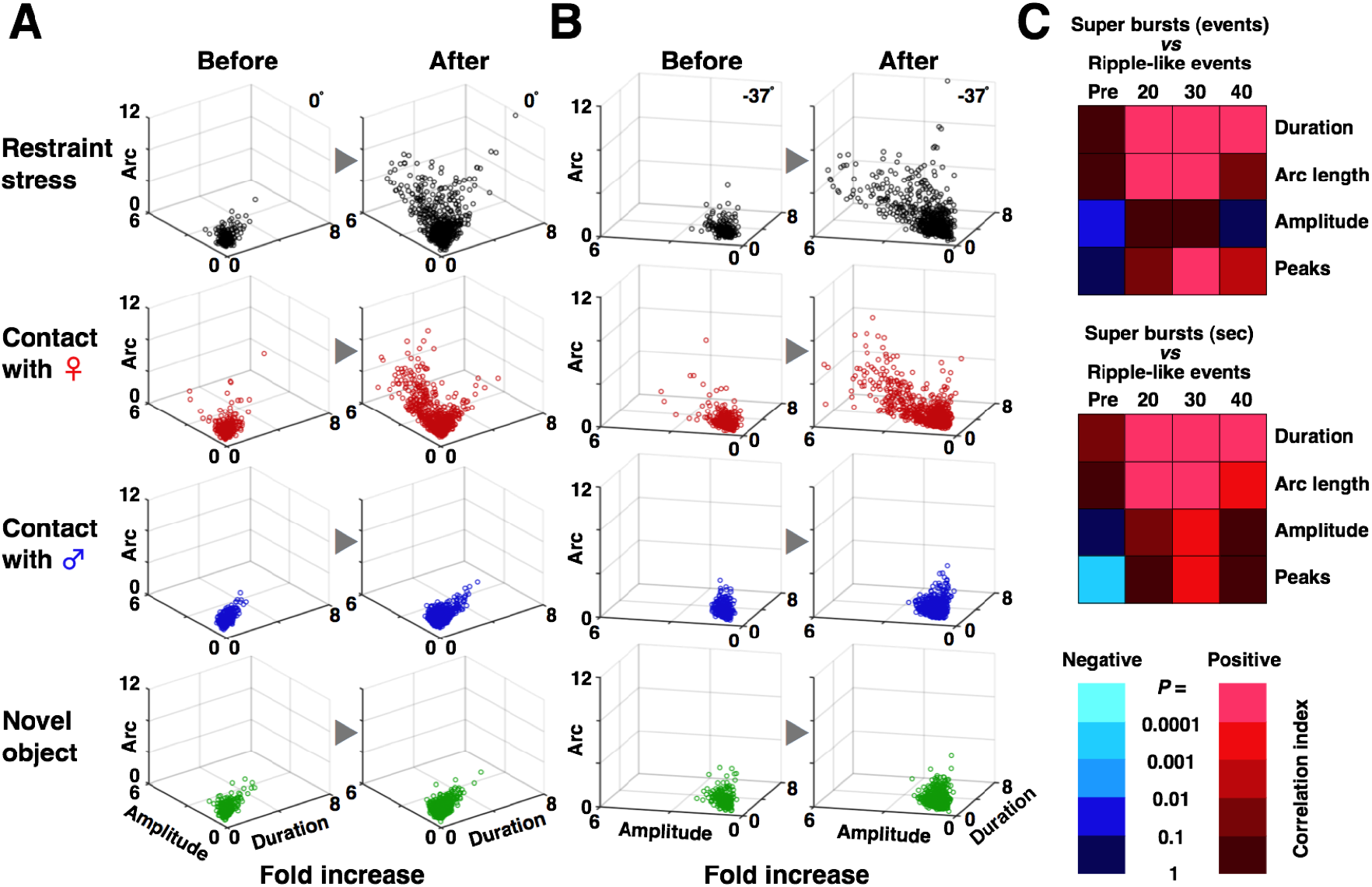
Plots of the features for individual ripple-like events. (A) Instead of fourdimensional plots, we plotted three-dimensionally the three features (amplitude, duration, arc length) of a single ripple-like event before (left) and after (right) the experiences. (B) Same plots in a different horizontal angle (37° rotate left). MANOVA in four-dimensional virtual plots revealed both experience-induced changes and experience-specificity (*P* < 0.0001). (C) The features of ripple-like events after the experience were positively correlated with the number (upper) or the duration of super bursts (middle). Heat map (lower) indicates the strength of the negative or positive correlation.

For amplitude analysis (Fig. 4B), we calculated the difference between the lowest and highest peaks. Arc length (*L*: Fig. 4F) was calculated using the following equation:

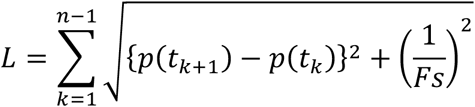

where *Fs* is the sampling rate for signal acquisition and *p*(*t*) indicates voltage value at time point *t*. The total number of sampling points during the duration of a ripple-like event is *n* (Fig. 4D). For peak analysis (Fig. 4H), we counted the number of negative peaks during the ripple-like firing events. For graphic expression (Figs. 5A and B), three of four parameters were multi-dimensionally plotted using MATLAB^®^ (The MathWorks, Inc. MA).

### Experimental schedule

Rats were handled for at least one week before surgery and allowed to recover from surgery for at least 2 weeks before the recording experiments. On the day of the experiment, a recording electrode was inserted into the pyramidal cell layer of CA1 neurons. We started the recording of multiple-unit CA1 firings, and spontaneous behavior was simultaneously monitored while the animals were in their well-habituated home cages. Details of the recording schedule are described below. To confirm the formation of memory for each experience, behavioral responses were monitored again on the day following the first experience (Fig. 1B).

### Neural recording schedule

At least 15 min after the recording of basal condition, the rats were exposed to either the restraint stress or a first encounter with a female, male, or object for 10 min (Fig. 1A).

For restraint, rats were taken from the home cages, their legs were strapped using soft cotton ties, and the animals were fixed onto the wood board for 10 min (Mitsushima et al, 2006, 2008). The animals then were returned to their home cages, and the multiple-unit firings were sequentially recorded for more than 30 min. For the other experiences, a sexually mature female (postnatal age, 8–12 weeks), young male (postnatal age, 6–7 weeks), or novel object [yellow LEGO^®^/DUPLO^®^ brick, 15 cm (h) × 8 cm (w) × 3 cm (d)] were placed in the home cage for 10 min. The bricks have a weak plastic smell and were fixed onto the side wall using double-sided tape. After the removal of the intruder or object, multiple-unit firings were sequentially recorded for more than 30 min.

### Histology

At the end of experiments, animals were deeply anesthetized with sodium pentobarbital (400 mg/kg, i.p.) and immediately perfused transcardially with a solution of 0.1 M phosphate buffer containing 4% paraformaldehyde. The brain was removed and then post-fixed with the same paraformaldehyde solution and immersed in 10%–30% sucrose solution. Coronal sections (40 μm thick) were stained with hematoxylin and eosin. The locations of cannulas, recoding electrode tips, and tracks in the brain were identified with the aid of a stereotaxic atlas (Paxinos & Watson, 2013).

### Slice-patch clamp analysis

Forty minutes after onset of the episodic experience, we anesthetized the rats with pentobarbital and prepared acute brain slices (Mitsushima et al, 2011, 2013). The inexperienced control rats were injected with the same dose of anesthesia in their home cage. Whole-cell recordings were performed as described previously (Kida et al, 2016). Briefly, the brains were quickly perfused with ice-cold dissection buffer (25.0 mM NaHCO_3_, 1.25 mM NaH_2_PO_4_, 2.5 mM KCl, 0.5 mM CaCl_2_, 7.0 mM MgCl_2_, 25.0 mM glucose, 90 mM choline chloride, 11.6 mM ascorbic acid, 3.1 mM pyruvic acid) and gassed with 5% CO_2_ / 95% O_2_. Coronal brain slices (target CA1 area: anteroposterior - 3.8 mm, dorsal-ventral 2.5 mm, mediolateral ± 2.0 mm) were cut (350 μm, Leica vibratome, VT-1200) in dissection buffer and transferred to physiological solution (22– 25°C, 114.6 mM NaCl, 2.5 mM KCl, 26 mM NaHCO_3_, 1 mM NaH_2_PO_4_, 10 mM glucose, 4 mM MgCl_2_, 4 mM CaCl_2_, pH 7.4, gassed with 5% CO_2_ / 95% O_2_). The recording chamber was perfused with physiological solution containing 0.1 mM picrotoxin and 4 μM 2-chloroadenosine at 22–25°C. For the miniature excitatory synaptic current (mEPSC) and miniature inhibitory synaptic current (mIPSC) recordings, we used a physiological solution containing 0.5 μM tetrodotoxin to block Na^+^ channels.

Patch recording pipettes (4–7 MΩ) were filled with intracellular solution (127.5 mM cesium methanesulfonate, 7.5 mM CsCl, 10 mM HEPES, 2.5 mM MgCl_2_, 4 mM Na_2_ATP, 0.4 mM Na_3_GTP, 10 mM sodium phosphocreatine, 0.6 mM EGTA at pH 7.25). Whole-cell recordings were obtained from CA1 pyramidal neurons from the rat hippocampus using an Axopatch 700A amplifier (Axon Instruments).

For the miniature recordings, the mEPSCs (−60 mV holding potential) and mIPSCs (0 mV holding potential) were recorded sequentially for 5 min in the same CA1 neuron (Fig. 6A). Bath application of an AMPA receptor blocker (CNQX, 10 μM) or GABA_A_ receptor blocker (bicuculline methiodide, 10 μM) consistently blocked the mEPSC or mIPSC events, respectively.

**Figure 6.**
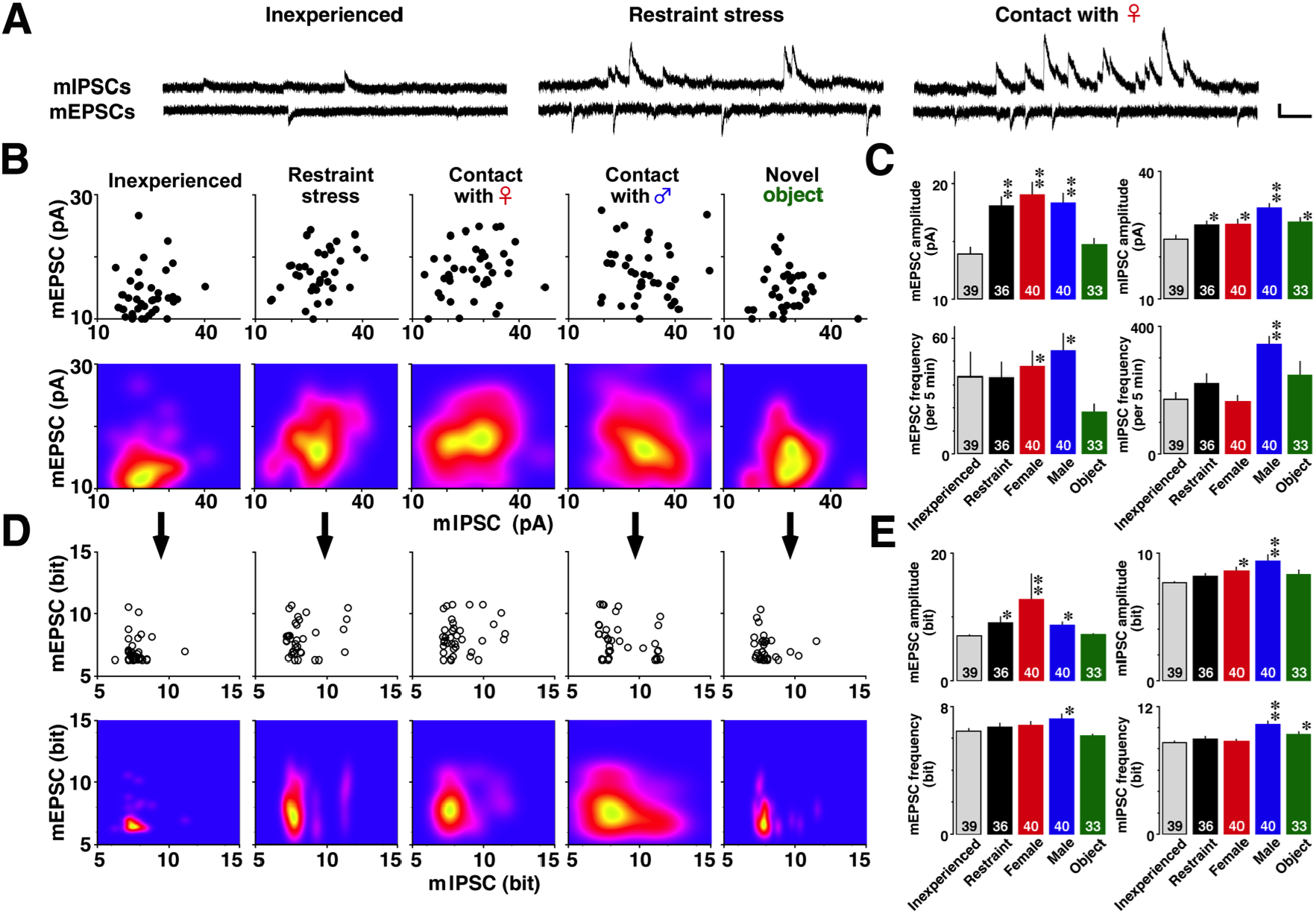
Experience-specific plasticity at excitatory and inhibitory CA1 synapses. (A) Representative traces of mEPSCs and mIPSCs. We measured mEPSCs at −60 mV and mIPSCs at 0 mV sequentially in the same pyramidal neuron to obtain four measures (amplitudes and frequencies of mEPSCs and mIPSCs). Vertical bar, 20 pA; horizontal bar, 200 ms. (B) Instead of four-dimensional plots, we two-dimensionally plotted mean mEPSC and mIPSC amplitudes in individual CA1 neurons (upper) and visualized the density distribution (lower). MANOVA in four-dimensional virtual plots showed experience-specific synaptic plasticity (*P* < 0.0001). (C) Differences in the individual four parameters (one-way ANOVA). (D) Based on the appearance probability in the inexperienced group, we further calculated self-entropy in individual CA1 neurons. MANOVA in four-dimensional virtual plots showed experience-specific distribution of self-entropy parameters (*P* < 0.0001). (E) Difference in the individual four self-entropy parameters (one-way ANOVA). The number of cells in each group is shown at the bottom of each bar. \**P* < 0.05, \*\**P* < 0.01 *vs*. inexperienced. Error bars indicate ± SEM.

### Self-entropy analysis

We calculated the self-entropy per neuron as reported previously (Sakimoto et al, 2019). First, we determined the distribution of appearance probability of mean mEPSC and mIPSC amplitudes separately using one-dimensional kernel density analysis. Geometric/topographic features of the appearance probability were calculated using kernel density analysis, as follows: Let *X*_1_, *X*_2_,…, *X_n_* denote a sample of size *n* from real observations. The kernel density estimate of *P* at the point *x* is given by

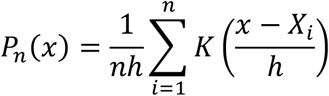

where *K* is a smooth function called the Gaussian kernel function and *h* > 0 is the smoothing bandwidth that controls the amount of smoothing. We chose Silverman’s reference bandwidth or Silverman’s rule of thumb (Sheather, 2004, Silverman, 1986). It is given by

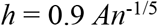

where *A* = min [sample SD, interquartile range/1.34]. For graphic expression, the distribution was visualized two dimensionally (Figs. 6B and 6D) in the *R* software environment (*R* Foundation for Statistical Computing, Vienna, Austria). By normalizing integral values in inexperienced controls, we found the distribution of appearance probability at any point. Then, we calculated the appearance probability at selected points. All data points for probability in inexperienced and experienced rats were converted to self-entropy (bit) using the Shannon entropy concept, defined from the Information Theory (Shannon, 1948).

### Statistical analysis

Data and statistical analyses were performed using SPSS or StatView software. To analyze behavioral parameters, we used paired *t*-tests. To compare temporal dynamics in the duration of the silent period or number of ripple-like events, we used the Friedman test followed by the *post hoc* Wilcoxon signed-rank test. Differences in the ripple-like events or super bursts or silent period or features among experiences were compared using the Kruskal–Wallis test followed by the *post hoc* Mann–Whitney *U* test. We applied Pearson’s correlation coefficient to examine correlation among super bursts, ripple-like events, and silent period.

The number and total duration of super bursts or features of ripple-like events, such as amplitude, duration, arc length, and peak, were analyzed using two-way analysis of variance (ANOVA) with the between-group factors of time and experience, followed by *post hoc* ANOVAs and Fisher’s *post hoc* least significance difference test. Each miniature parameter and each self-entropy value were analyzed using one-way factorial ANOVA in which the experience was the between-group factor.

Overall differences in two features of super bursts, four features of individual ripple-like events, or four miniature parameters (mEPSC and mIPSC amplitudes and frequencies) were analyzed using multivariate ANOVA (MANOVA) with Wilks’ Lambda distribution. Specific differences between two experiences were further analyzed using *post hoc* MANOVAs. The Shapiro-Wilk test and *F*-test were used to determine normality and equality of variance, respectively. Because the data had large variations within a group, we performed *log* (*1+x*) transformation prior to the analysis (Mitsushima et al, 1994). *P* < 0.05 was considered statistically significant.

## Results

### Evaluation of experienced memory

The male rats being recorded vigorously resisted and vocalized frequently during the 10 min of restraint (Mitsushima et al, 2006). The males checked, chased, and sometimes mounted a female intruder, but no intromission or ejaculation was observed (Mitsushima et al, 2009). With a male intruder, the males attacked him minutes after the check and hesitation. With the novel object, the animals checked it and sometimes smelled and baited it.

To evaluate the acquired memory, we compared behavioral parameters during the first experience with those during a second experience (Fig. 1B). The rats that experienced the 10-min restraint stress showed a smaller number of vocalizations in the second exposure (*t_6_* = 3.476, *P* = 0.0129). Similarly, those that encountered a female, male, or novel object consistently in the second encounter showed a shortened latency to vaginal inspection (*t_8_* = 3.492, *P* = 0.0082) and attack (*t_7_* = 4.192, *P* = 0.0041) and a shorter observation time (*t_9_* = 2.901, *P* = 0.0176), suggesting acquisition of experienced memory.

### Multiple-unit firings

To record multiple-unit firings of CA1 neurons, we implanted a depth-adjustable tungsten electrode with a resistance of 50 to 80 kΩ (Fig. 1D). We extracted super bursts (Fig. 1G), silent periods (Fig. 1H), or ripple-like events (Fig. 1I) based on basal firing rate in their habituated home cage. Per the criteria, spontaneous super burst events showed a firing rate and silent periods showed a duration of inter-spike interval greater than 3 SD of basal firings. Ripple-like events were short-term high-frequency firings (≈ 50 ms) with more than a 1:6 signal-to-noise ratio. Although spike sorting was not successful in super bursts and ripple-like events, coordinated firings of multiple single neurons formed a single event of super burst or ripple-like firings (Fig. 1E). Single events of a silent period lasted 0.89 ± 0.04 s (*N* = 278).

### Super bursts

Before the test experience, the rats in their habituated home cage mainly exhibited sporadic spontaneous firings. After the onset of an experience, they frequently showed super burst events (Fig. 2A). Both the incidence [Fig. 2C (Table 1)] and duration of super bursts [Fig. 2D (Table 2)] changed dramatically from baseline to the experience. To analyze the super bursts, we used two-way ANOVA with experience as the between-group factor and time as the within-group factor. The results showed a significant main effect of experience (events 3 min^-1^, *F*_3, 403_ = 4.461, *P* = 0.010; duration, *F*_3, 403_ = 5.434, *P* = 0.004), time (events 3 min^-1^, *F*_13, 403_ = 10.194, *P* < 0.0001; duration, *F*_13, 403_ = 14.061, *P* < 0.0001), and their interaction (events 3 min^-1^, *F*_36, 403_ = 2.368, *P* < 0.0001; duration, *F*_39, 403_ = 4.626, *P* < 0.0001).

**Table 1.** *P* values for *post hoc* analysis of Figure 2C

|  | Min after the onset |  |  |  |  |  |  |  |  |  |  |  |  |
| --- | --- | --- | --- | --- | --- | --- | --- | --- | --- | --- | --- | --- | --- |
|  | 3 | 6 | 9 | 12 | 15 | 18 | 21 | 24 | 27 | 30 | 33 | 36 | 39 |
| **Restraint** | 0.003 | 0.478 | 0.373 | 0.738 | 0.966 | 0.216 | 0.216 | 0.216 | 0.216 | 0.216 | 0.216 | 0.216 | 0.216 |
| **Female** | 0.134 | 0.058 | 0.422 | 0.012 | 0.316 | 0.244 | 0.460 | 0.327 | 0.670 | 0.389 | 0.018 | 0.018 | 0.018 |
| **Male** | 0.057 | 0.383 | 0.802 | 0.504 | 0.427 | 0.298 | 0.423 | 0.150 | 0.005 | 0.927 | 0.150 | 0.005 | 0.423 |
| **Object** | 0.465 | 0.824 | 0.465 | 0.033 | 0.033 | 0.465 | 0.033 | 0.033 | 0.033 | 0.033 | 0.033 | 0.033 | 0.033 |

**Table 2.** *P* values for *post hoc* analysis of Figure 2D

|  | Min after the onset |  |  |  |  |  |  |  |  |  |  |  |  |
| --- | --- | --- | --- | --- | --- | --- | --- | --- | --- | --- | --- | --- | --- |
|  | 3 | 6 | 9 | 12 | 15 | 18 | 21 | 24 | 27 | 30 | 33 | 36 | 39 |
| **Restraint** | 0.001 | 0.483 | 0.407 | 0.947 | 0.549 | 0.218 | 0.218 | 0.218 | 0.218 | 0.218 | 0.218 | 0.218 | 0.218 |
| **Female** | 0.144 | 0.060 | 0.416 | 0.010 | 0.400 | 0.325 | 0.583 | 0.384 | 0.057 | 0.454 | 0.020 | 0.020 | 0.020 |
| **Male** | 0.021 | 0.145 | 0.342 | 0.408 | 0.884 | 0.912 | 0.646 | 0.145 | 0.035 | 0.316 | 0.623 | 0.035 | 0.904 |
| **Object** | 0.214 | 0.612 | 0.357 | 0.058 | 0.058 | 0.498 | 0.058 | 0.058 | 0.058 | 0.058 | 0.058 | 0.058 | 0.058 |

To further analyze the main effect of time in individual experiences, we performed *post hoc* ANOVAs. Both incidence and duration of super bursts were significantly increased by restraint stress (events 3 min^-1^, *F*_13, 91_ = 6.265, *P* < 0.0001; duration, *F*_13, 91_ = 9.251, *P* < 0.0001), contact with a female (events 3 min^-1^, *F*_13, 91_ = 3.408, *P* = 0.0003; duration, *F*_13, 91_ = 4.166, *P* < 0.0001), and contact with a male (events 3 min^-1^, *F*_13, 117_ = 2.630, *P* = 0.003; duration, *F*_13, 117_ = 2.894, *P* = 0.0012). They remained unchanged after contact with a novel object (events 3 min^-1^, *F*_13, 104_ = 1.684, *P* = 0.075; duration, *F*_13, 104_ = 1.759, *P* = 0.059). For the between-groups comparison, *post hoc* analysis showed specific differences in incidence [Fig. 2E left (Table 3)] or their duration [Fig. 2E right (Table 4)].

**Table 3.** *P* values for *post hoc* analysis of Figure 2E left

|  | Before | Min after the onset |  |  |  |  |  |  |  |  |  |  |  |  |
| --- | --- | --- | --- | --- | --- | --- | --- | --- | --- | --- | --- | --- | --- | --- |
|  |  | 3 | 6 | 9 | 12 | 15 | 18 | 21 | 24 | 27 | 30 | 33 | 36 | 39 |
| **Restraint vs Female** | 0.183 | 0.242 | 0.151 | 0.808 | 0.001 | 0.072 | 0.014 | 0.031 | 0.009 | 0.147 | 0.579 | - | - | - |
| **Restraint vs Male** | 0.133 | 0.073 | 0.969 | 0.638 | 0.390 | 0.909 | 0.538 | 0.451 | 0.704 | - | 0.126 | 0.225 | - | 0.073 |
| **Restraint vs Object** | 0.356 | 0.002 | 0.638 | 0.306 | 0.600 | 0.679 | 0.673 | - | - | - | - | - | - | - |
| **Female vs Male** | 0.916 | 0.552 | 0.121 | 0.469 | 0.004 | 0.074 | 0.042 | 0.115 | 0.015 | 0.127 | 0.335 | 0.225 | - | 0.073 |
| **Female vs Object** | 0.646 | 0.040 | 0.056 | 0.205 | 0.000 | 0.026 | 0.032 | 0.027 | 0.007 | 0.136 | 0.568 |  | - | - |
| **Male vs Object** | 0.552 | 0.109 | 0.652 | 0.546 | 0.154 | 0.579 | 0.849 | 0.437 | 0.694 | - | 0.115 | 0.211 | - | 0.065 |

**Table 4.** *P* values for *post hoc* analysis of Figure 2E right

|  | Before | Min after the onset |  |  |  |  |  |  |  |  |  |  |  |  |
| --- | --- | --- | --- | --- | --- | --- | --- | --- | --- | --- | --- | --- | --- | --- |
|  |  | 3 | 6 | 9 | 12 | 15 | 18 | 21 | 24 | 27 | 30 | 33 | 36 | 39 |
| **Restraint vs Female** | 0.364 | 0.003 | 0.357 | 0.925 | 0.000 | 0.046 | 0.020 | 0.027 | 0.006 | 0.147 | 0.184 | - | - | - |
| **Restraint vs Male** | 0.691 | < 0.001 | 0.804 | 0.365 | 0.697 | 0.896 | 0.463 | 0.353 | 0.901 | - | 0.617 | 0.225 | - | 0.073 |
| **Restraint vs Object** | 0.906 | < 0.001 | 0.454 | 0.177 | 0.610 | 0.780 | 0.691 | - | - | - | - | - | - | - |
| **Female vs Male** | 0.150 | 0.347 | 0.226 | 0.419 | < 0.001 | 0.048 | 0.076 | 0.142 | 0.005 | 0.127 | 0.362 | 0.225 | - | 0.073 |
| **Female vs Object** | 0.295 | 0.012 | 0.095 | 0.209 | < 0.001 | 0.022 | 0.043 | 0.023 | 0.005 | 0.136 | 0.172 | - | - | - |
| **Male vs Object** | 0.775 | 0.229 | 0.590 | 0.613 | 0.349 | 0.668 | 0.734 | 0.338 | 0.898 | - | 0.605 | 0.211 | - | 0.065 |

Both events and their duration were two-dimensionally compared using repeated measures of MANOVA (Fig. 2E), with experience as the between-group factor and time as the within-group factor. The main effect of experience (*F*_6, 970_ = 11.408, *P* < 0.0001) and the *post hoc* analyses indicated specific differences between two experiences, suggesting episode-specific temporal dynamics of the super bursts (Table 5).

**Table 5.** MANOVA and *post hocs* for Figure 2E

|  | Super bursts | Silent periods |
| --- | --- | --- |
| **Overall** | $F_{6,970} = 11.408, P < 0.001^{**}$ | $F_{6,210} = 1.566, P = 0.158$ |
| **Restraint vs Female** | $F_{2,221} = 14.389, P < 0.001^{**}$ | $F_{2,50} = 1.929, P = 0.156$ |
| **Restraint vs Male** | $F_{2,249} = 10.967, P < 0.001^{**}$ | $F_{2,57} = 2.492, P = 0.092$ |
| **Restraint vs Object** | $F_{2,235} = 6.023, P = 0.003^{**}$ | $F_{2,54} = 0.302, P = 0.741$ |
| **Female vs Male** | $F_{2,249} = 7.891, P < 0.001^{**}$ | $F_{2,50} = 0.006, P = 0.994$ |
| **Female vs Object** | $F_{2,235} = 16.638, P < 0.001^{**}$ | $F_{2,47} = 2.178, P = 0.125$ |
| **Male vs Object** | $F_{2,263} = 7.281, P < 0.001^{**}$ | $F_{2,54} = 2.496, P = 0.092$ |

The duration and relative firing rate of individual super bursts were further analyzed and plotted two-dimensionally (triangles in Fig. 2F). X-axis represents duration (s), and Y-axis represents times over the threshold (+3 SD of the mean firing rate before the experience). The two features were compared using MANOVA, with experience as the between-group factor. The results indicated a significant main effect of experience (*F*_6, 554_ = 29.201, *P* < 0.0001). *Post hoc* MANOVA showed specific differences between two experiences, suggesting episode-specific features of individual super bursts (Table 6).

**Table 6.**
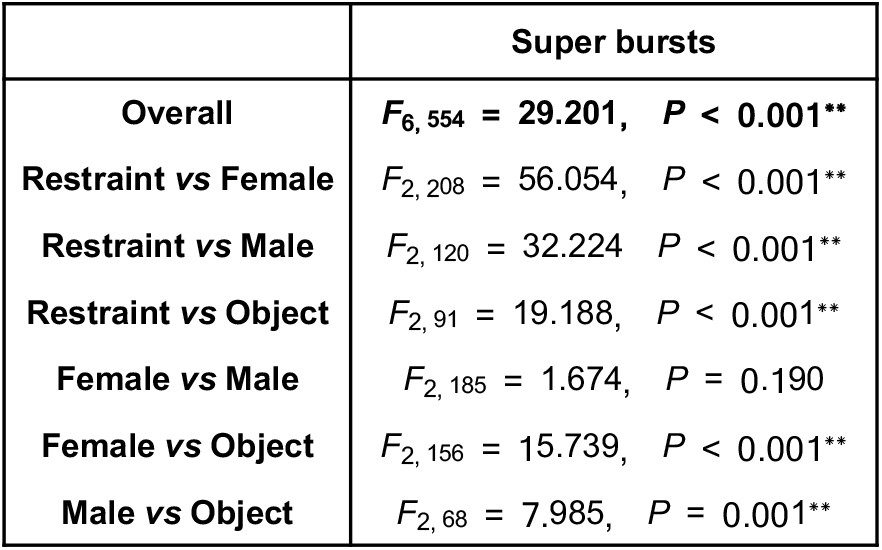
MANOVA and *post hocs* for Figure 2F

### Silent periods

We found a within-group temporal change in the duration of silent periods with restraint stress (Fig. 3B; *χ*^2^_2_ = 6.200, *P* = 0.045) and female (*χ*^2^_2_ = 7.000, *P* = 0.030) and male contact (*χ*^2^_2_ = 4.200, *P* = 0.123). *Post hoc* tests further demonstrated a temporal difference in the total duration, which was significantly increased 20 min after the onset of restraint stress (*Z* = −2.090, *P* = 0.037) and 30 min after the onset of female (*Z* = −2.100, *P* = 0.036) and male contact (*Z* = −2.395, *P* = 0.017). The novel object was associated with increased total duration (*χ*^2^_2_ = 2.889, *P* = 0.235). In the comparison between groups, we observed no significant difference in total duration (Fig. 3C; Kruskal–Wallis test; 20 min, *H*_3_ = 1.451 *P* = 0.693; 40 min, *H*_3_ = 0.588, *P* = 0.899).

To examine episode specificity, we compared both incidence and duration two-dimensionally using MANOVA, with experience as the between-group factor and time as the within-group factor. MANOVA results showed no significant main effect of experience (*F*_6, 210_ = 1.566, *P* = 0.158), suggesting that the temporal dynamics of silent periods may not be episode-specific (Table 5).

### Ripple-like events

We found a significant temporal change in the incidence of ripple-like events within the group for restraint stress (*χ*^2^_2_ = 9.805, *P* = 0.007) and female contact (*χ*^2^_2_ = 7.000, *P* = 0.030), but not for contact with a male (*χ*^2^_2_ = 0.200, *P* = 0.905) or novel object (*χ*^2^_2_ = 2.667, *P* = 0.264; Fig. 3F). The incidence significantly increased 20 min after the onset of restraint stress (*Z* = −2.805, *P* = 0.005) and 40 min after the onset of female contact (*Z* = −2.240, *P* = 0.025).

In the comparison between groups, the *post hoc* test demonstrated a significant main effect within 20 min (*H*_3_ = 10.439, *P* = 0.015) and 40 min (*H*_3_ = 9.181, *P* = 0.027) after the onset of experience. Figure 3G (Table 7) shows detailed differences among experiences.

**Table 7.** *Post hoc* analysis for Figure 3G

|  | Min after the onset |  |
| --- | --- | --- |
|  | 20 | 40 |
| **Restraint vs Female** | $Z = -1.599, P = 0.110$ | $Z = -0.089, P = 0.929$ |
| **Restraint vs Male** | $Z = -2.570, P = 0.010$ | $Z = -1.081, P = 0.070$ |
| **Restraint vs Object** | $Z = -2.531, P = 0.011$ | $Z = -2.205, P = 0.028$ |
| **Female vs Male** | $Z = -1.333, P = 0.183$ | $Z = -1.866, P = 0.062$ |
| **Female vs Object** | $Z = -1.540, P = 0.124$ | $Z = -2.406, P = 0.016$ |
| **Male vs Object** | $Z = -0.816, P = 0.414$ | $Z = -0.816, P = 0.414$ |

### Correlation of super bursts with ripple-like events or silent periods

Both the incidence and duration of super bursts were significantly correlated with the relative number of ripple-like events (Figs. 3H and 3I) but not with silent periods (Figs. 3D and 3E). The results suggest a role for super bursts in increasing ripple-like events. Detailed correlations of multiple features with ripple-like events are shown in Figure 5C.

### Features of individual ripple-like events

We extracted four features of individual ripple-like events (Fig. 4A) and analyzed them using two-way ANOVA, with experience as the between-group factor and time as the within-group factor (Table 8). Further *post hoc* ANOVAs and the temporal dynamics are shown for amplitude in Figure 4B (Table 9), for duration in Figure 4D (Table 10), for arc length in Figure 4F (Table 11), and for peaks in Figure 4H (Table 12). Values for all of these features significantly increased with restraint stress and female contact. In contrast, only a partial and different effect was seen with contact with a male or object. Duration and arc length increased with contact with a male, but amplitude and peak number did not. The number of peaks decreased with contact with a novel object, but other features did not change. Differences between two specific groups are shown for amplitude in Figure 4C (Table 13), for duration in Figure 4E (Table 14), for arc length in Figure 4G (Table 15), and for peaks in Figure 4I (Table 16).

**Table 8.** Two-way ANOVA for individual features of ripple-like events

| Features | Experience | Time | Interaction |
| --- | --- | --- | --- |
| Amplitude | $F_{3, 4060} = 71.633, P < 0.001^{**}$ | $F_{3, 4060} = 24.094, P < 0.001^{**}$ | $F_{9, 4060} = 8.414, P < 0.001^{**}$ |
| Duration | $F_{3, 4060} = 45.261, P < 0.001^{**}$ | $F_{3, 4060} = 31.951, P < 0.001^{**}$ | $F_{9, 4060} = 5.994, P < 0.001^{**}$ |
| Arc length | $F_{3, 4060} = 91.713, P < 0.001^{**}$ | $F_{3, 4060} = 40.587, P < 0.001^{**}$ | $F_{9, 4060} = 10.840, P < 0.001^{**}$ |
| Peaks | $F_{3, 4060} = 93.602, P < 0.001^{**}$ | $F_{3, 4060} = 21.460, P < 0.001^{**}$ | $F_{9, 4060} = 11.739, P < 0.001^{**}$ |

**Table 9.** *Post hoc* analysis for Figure 4B

|  | Overall | Min after the onset |  |  |
| --- | --- | --- | --- | --- |
|  |  | 20 | 30 | 40 |
| Restraint | $F_{3, 962} = 17.699, P < 0.001^{**}$ | $P < 0.001^{**}$ | $P < 0.001^{**}$ | $P < 0.001^{**}$ |
| Female | $F_{3, 1076} = 15.533, P < 0.001^{**}$ | $P < 0.001^{**}$ | $P < 0.001^{**}$ | $P < 0.001^{**}$ |
| Male | $F_{3, 1070} = 0.626, P = 0.598$ | $P = 0.256$ | $P = 0.638$ | $P = 0.898$ |
| Object | $F_{3, 952} = 0.107, P = 0.956$ | $P = 0.610$ | $P = 0.702$ | $P = 0.896$ |

**Table 10.** *Post hoc* analysis for Figure 4D

|  | Overall | Min after the onset |  |  |
| --- | --- | --- | --- | --- |
|  |  | 20 | 30 | 40 |
| Restraint | $F_{3, 962} = 23.385, P < 0.001^{**}$ | $P < 0.001^{**}$ | $P < 0.001^{**}$ | $P < 0.001^{**}$ |
| Female | $F_{3, 1076} = 7.362, P < 0.001^{**}$ | $P < 0.001^{**}$ | $P = 0.002^{**}$ | $P < 0.001^{**}$ |
| Male | $F_{3, 1070} = 10.165, P < 0.001^{**}$ | $P = 0.041^*$ | $P < 0.001^{**}$ | $P < 0.001^{**}$ |
| Object | $F_{3, 952} = 0.552, P = 0.647$ | $P = 0.441$ | $P = 0.961$ | $P = 0.293$ |

**Table 11.** *Post hoc* analysis for Figure 4F

|  | Overall | Min after the onset |  |  |
| --- | --- | --- | --- | --- |
|  |  | 20 | 30 | 40 |
| Restraint | $F_{3, 962} = 25.564, P < 0.001^{**}$ | $P < 0.001^{**}$ | $P < 0.001^{**}$ | $P < 0.001^{**}$ |
| Female | $F_{3, 1076} = 19.331, P < 0.001^{**}$ | $P < 0.001^{**}$ | $P < 0.001^{**}$ | $P < 0.001^{**}$ |
| Male | $F_{3, 1070} = 4.080, P = 0.007^{**}$ | $P = 0.674$ | $P = 0.005^{**}$ | $P = 0.011^*$ |
| Object | $F_{3, 952} = 0.109, P = 0.955$ | $P = 0.780$ | $P = 0.766$ | $P = 0.990$ |

**Table 12.** *Post hoc* analysis for Figure 4H

|  | Overall | Min after the onset |  |  |
| --- | --- | --- | --- | --- |
|  |  | 20 | 30 | 40 |
| **Restraint** | $F_{3, 962} = 21.125, P < 0.001^{**}$ | $P < 0.001^{**}$ | $P < 0.001^{**}$ | $P < 0.001^{**}$ |
| **Female** | $F_{3, 1076} = 11.643, P < 0.001^{**}$ | $P < 0.001^{**}$ | $P < 0.001^{**}$ | $P < 0.001^{**}$ |
| **Male** | $F_{3, 1070} = 3.148, P = 0.024^*$ | $P = 0.206$ | $P = 0.002^{**}$ | $P = 0.087$ |
| **Object** | $F_{3, 952} = 8.215, P < 0.001^{**}$ | $P = 0.030^*$ | $P < 0.001^{**}$ | $P < 0.001^{**}$ |

**Table 13.** *Post hoc* analysis for Figure 4C

|  | Min after the onset |  |  |
| --- | --- | --- | --- |
|  | 20 | 30 | 40 |
| **Overall** | $F_{3, 990} = 24.955, P < 0.001$ | $F_{3, 1033} = 29.659, P < 0.001$ | $F_{3, 993} = 25.285, P < 0.001$ |
| **Restraint vs Female** | $P = 0.199$ | $P = 0.364$ | $P = 0.109$ |
| **Restraint vs Male** | $P < 0.001^{**}$ | $P < 0.001^{**}$ | $P < 0.001^{**}$ |
| **Restraint vs Object** | $P < 0.001^{**}$ | $P < 0.001^{**}$ | $P < 0.001^{**}$ |
| **Female vs Male** | $P < 0.001^{**}$ | $P < 0.001^{**}$ | $P < 0.001^{**}$ |
| **Female vs Object** | $P < 0.001^{**}$ | $P < 0.001^{**}$ | $P < 0.001^{**}$ |
| **Male vs Object** | $P = 0.309$ | $P = 0.609$ | $P = 0.976$ |

**Table 14.**
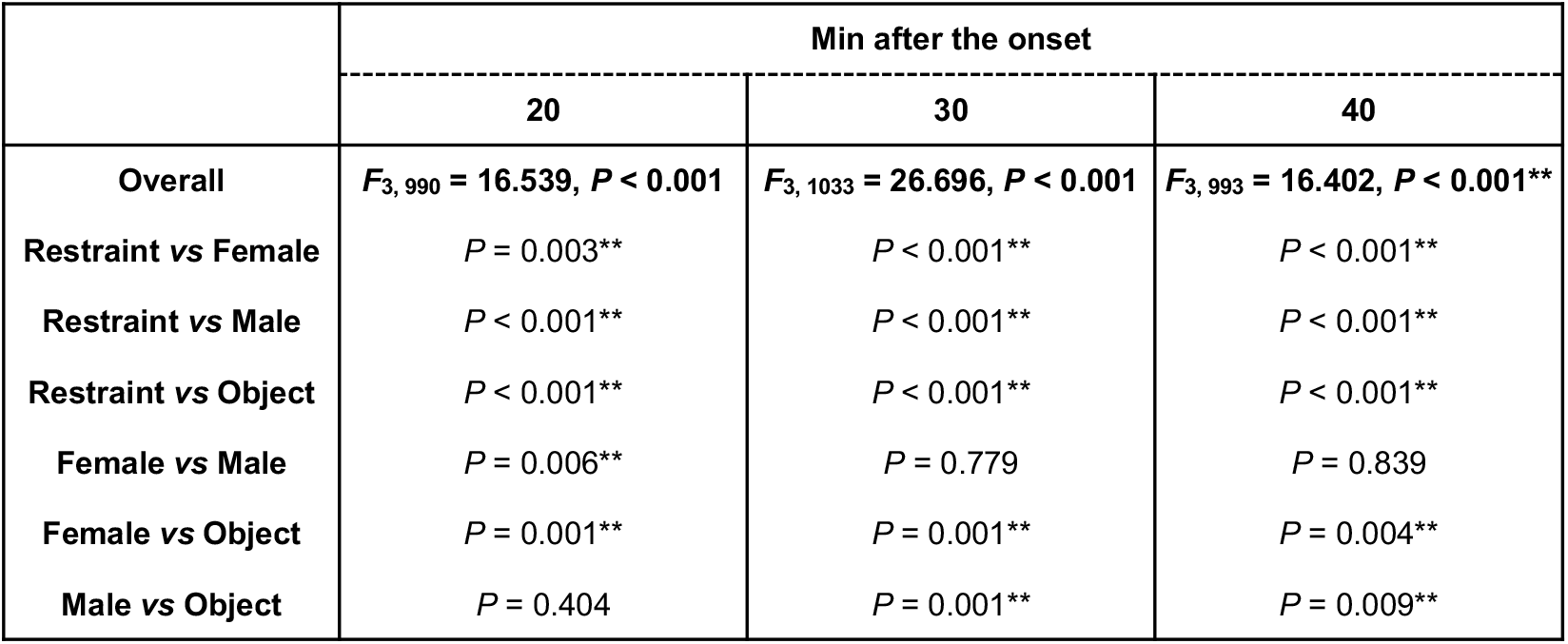
*Post hoc* analysis for Figure 4E

|  | Min after the onset |  |  |
| --- | --- | --- | --- |
|  | 20 | 30 | 40 |
| **Overall** | $F_{3, 990} = 16.539, P < 0.001$ | $F_{3, 1033} = 26.696, P < 0.001$ | $F_{3, 993} = 16.402, P < 0.001^{**}$ |
| **Restraint vs Female** | $P = 0.003^{**}$ | $P < 0.001^{**}$ | $P < 0.001^{**}$ |
| **Restraint vs Male** | $P < 0.001^{**}$ | $P < 0.001^{**}$ | $P < 0.001^{**}$ |
| **Restraint vs Object** | $P < 0.001^{**}$ | $P < 0.001^{**}$ | $P < 0.001^{**}$ |
| **Female vs Male** | $P = 0.006^{**}$ | $P = 0.779$ | $P = 0.839$ |
| **Female vs Object** | $P = 0.001^{**}$ | $P = 0.001^{**}$ | $P = 0.004^{**}$ |
| **Male vs Object** | $P = 0.404$ | $P = 0.001^{**}$ | $P = 0.009^{**}$ |

**Table 15.**
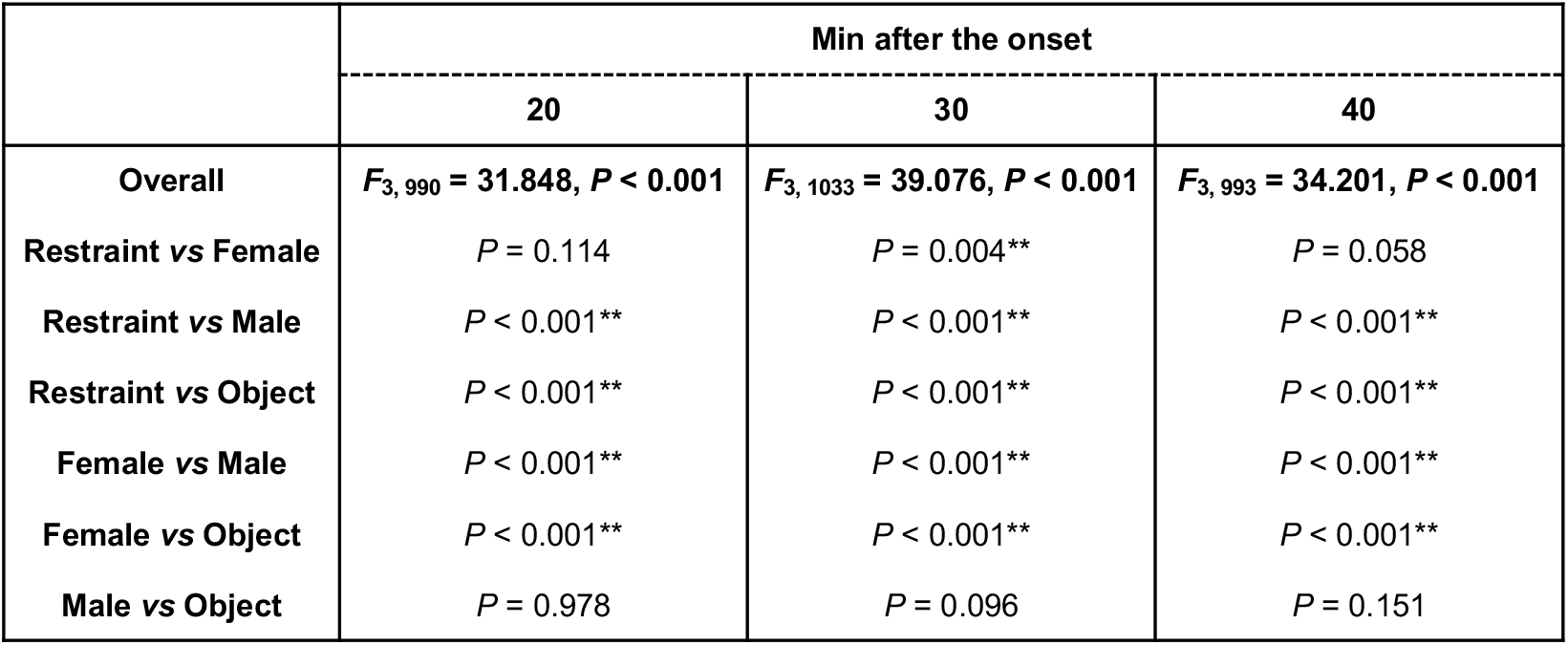
*Post-hoc* analysis for Figure 4G

**Table 16.** *Post hoc* analysis for Figure 4I

|  | Min after the onset |  |  |
| --- | --- | --- | --- |
|  | 20 | 30 | 40 |
| **Overall** | $F_{3, 990} = 34.445, P < 0.001^{**}$ | $F_{3, 1033} = 49.182, P < 0.001^{**}$ | $F_{3, 993} = 36.244, P < 0.001^{**}$ |
| **Restraint vs Female** | $P = 0.029^*$ | $P < 0.001^{**}$ | $P = 0.088$ |
| **Restraint vs Male** | $P < 0.001^{**}$ | $P < 0.001^{**}$ | $P < 0.001^{**}$ |
| **Restraint vs Object** | $P < 0.001^{**}$ | $P < 0.001^{**}$ | $P < 0.001^{**}$ |
| **Female vs Male** | $P < 0.001^{**}$ | $P = 0.026^*$ | $P < 0.001^{**}$ |
| **Female vs Object** | $P < 0.001^{**}$ | $P < 0.001^{**}$ | $P < 0.001^{**}$ |
| **Male vs Object** | $P = 0.008^{**}$ | $P < 0.001^{**}$ | $P = 0.001^{**}$ |

To examine further the diversified features of ripple-like events, we compared the four features using 2-way MANOVA, with experience and time as the between-group factors. Although a four-dimensional plot is quite difficult to depict, an example of three-dimensional plots of three features (amplitude, duration, and arc length) is shown in Figures 5A and B. The MANOVA results indicated a significant main effect of experience (*F*_12, 10734_ = 36.199, *P* < 0.0001), time (*F*_12, 10734_ = 12.825, *P* < 0.0001), and their interaction (*F*_36, 15205_ = 4.698, *P* < 0.0001). *Post hoc* MANOVA further suggested differences between two specific experiences (Table 17), suggesting episode-specific features of individual ripple-like events.

**Table 17.**
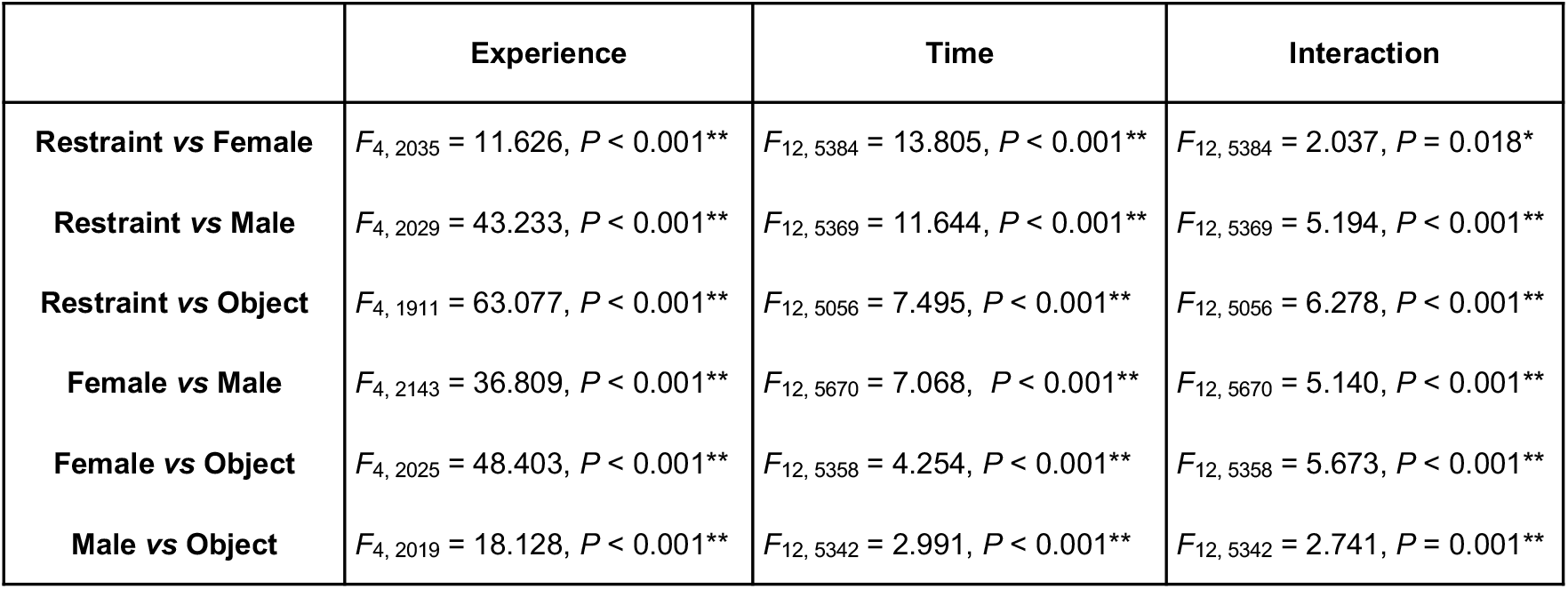
*Post hoc* MANOVAs for Figures 5A and 5B

|  | Experience | Time | Interaction |
| --- | --- | --- | --- |
| **Restraint vs Female** | $F_{4, 2035} = 11.626, P < 0.001^{**}$ | $F_{12, 5384} = 13.805, P < 0.001^{**}$ | $F_{12, 5384} = 2.037, P = 0.018^*$ |
| **Restraint vs Male** | $F_{4, 2029} = 43.233, P < 0.001^{**}$ | $F_{12, 5369} = 11.644, P < 0.001^{**}$ | $F_{12, 5369} = 5.194, P < 0.001^{**}$ |
| **Restraint vs Object** | $F_{4, 1911} = 63.077, P < 0.001^{**}$ | $F_{12, 5056} = 7.495, P < 0.001^{**}$ | $F_{12, 5056} = 6.278, P < 0.001^{**}$ |
| **Female vs Male** | $F_{4, 2143} = 36.809, P < 0.001^{**}$ | $F_{12, 5670} = 7.068, P < 0.001^{**}$ | $F_{12, 5670} = 5.140, P < 0.001^{**}$ |
| **Female vs Object** | $F_{4, 2025} = 48.403, P < 0.001^{**}$ | $F_{12, 5358} = 4.254, P < 0.001^{**}$ | $F_{12, 5358} = 5.673, P < 0.001^{**}$ |
| **Male vs Object** | $F_{4, 2019} = 18.128, P < 0.001^{**}$ | $F_{12, 5342} = 2.991, P < 0.001^{**}$ | $F_{12, 5342} = 2.741, P = 0.001^{**}$ |

We also analyzed the main effect of time in individual experiences (Figs. 5A and B). The distribution of the four features of individual ripple-like events was significantly changed after the experience of restraint (*F*_12, 2538_ = 9.118, *P* < 0.0001), contact with a female (*F*_12, 2839_ = 6.065, *P* < 0.0001), contact with a male (*F*_12, 2823_ = 3.453, *P* < 0.0001), and contact with a novel object (*F*_12, 2511_ = 3.188, *P* = 0.0002). The results suggest experience-specific diversity of individual ripple-like events.

### Synaptic plasticity

To further examine experience-induced synaptic plasticity, we prepared *ex vivo* brain slices 30 min after the experiences (Fig. 6A). With sequential recording of mEPSCs (at −60 mV) and mIPSCs (at 0 mV) from the same neuron (Mitsushima et al, 2013), we measured four parameters from individual CA1 neurons: amplitudes and frequencies for both mEPSCs and mIPSCs.

Figure 6B shows cell-specific plots of the means of AMPA receptor–mediated excitatory currents *vs*. GABA_A_ receptor–mediated inhibitory currents. We used kernel analysis to visualize two-dimensionally the distribution of appearance probability in lower panels. Although inexperienced rats exhibited a low and narrow distribution range, experience diversified the input strength. The results of one-way ANOVA in individual parameters are shown in Figure 6C (Table 18).

**Table 18.** One-way ANOVAs and *post hocs* for Figure 6C

|  | mEPSC amplitude | mIPSC amplitude |
| --- | --- | --- |
| **Overall** | $F_{4,183} = 9.442, P < 0.001^{**}$ | $F_{4,183} = 5.227, P < 0.001^{**}$ |
| Inexperienced vs Restraint | $P < 0.001^{**}$ | $P = 0.043^*$ |
| Inexperienced vs Female | $P < 0.001^{**}$ | $P = 0.049^*$ |
| Inexperienced vs Male | $P < 0.001^{**}$ | $P < 0.001^{**}$ |
| Inexperienced vs Object | $P = 0.370$ | $P = 0.012^{**}$ |
| Restraint vs Female | $P = 0.568$ | $P = 0.904$ |
| Restraint vs Male | $P = 0.919$ | $P = 0.019^*$ |
| Restraint vs Object | $P = 0.003^{**}$ | $P = 0.586$ |
| Female vs Male | $P = 0.630$ | $P = 0.011^*$ |
| Female vs Object | $P < 0.001^{**}$ | $P = 0.500$ |
| Male vs Object | $P = 0.001^{**}$ | $P = 0.081$ |
|  | mEPSC frequency | mIPSC frequency |
| **Overall** | $F_{4,183} = 2.869, P = 0.025^*$ | $F_{4,183} = 7.955, P < 0.001^{**}$ |
| Inexperienced vs Restraint | $P = 0.069$ | $P = 0.274$ |
| Inexperienced vs Female | $P = 0.022^*$ | $P = 0.703$ |
| Inexperienced vs Male | $P = 0.024^*$ | $P < 0.001^{**}$ |
| Inexperienced vs Object | $P = 0.887$ | $P = 0.464$ |
| Restraint vs Female | $P = 0.674$ | $P = 0.141$ |
| Restraint vs Male | $P = 0.689$ | $P < 0.001^{**}$ |
| Restraint vs Object | $P = 0.060^{**}$ | $P = 0.740$ |
| Female vs Male | $P = 0.983$ | $P < 0.001^{**}$ |
| Female vs Object | $P = 0.020^*$ | $P = 0.271$ |
| Male vs Object | $P = 0.021^*$ | $P < 0.001^{**}$ |

To compare experience-induced plasticity, we plotted the four parameters in a four-dimensional virtual space to analyze both amplitude and frequency of mEPSC and mIPSC events in individual CA1 neurons. We used one-way MANOVA with experience as the between-group factor, which showed a significant main effect of experience (Fig. 6C; *F*_16, 551_ = 4.729, *P* < 0.001). *Post hoc* MANOVA further showed multiple differences between two specific experiences (Table 19), suggesting experience-specific synaptic plasticity.

**Table 19.** *Post hoc* MANOVAs for Figure 6C

| Group | Inexperienced | Restraint stress | Contact w/ female | Contact w/male | Novel object |
| --- | --- | --- | --- | --- | --- |
| Inexperienced | -- | $F_{4,70} = 5.305$ | $F_{4,74} = 5.729$ | $F_{4,74} = 15.762$ | $F_{4,67} = 3.767$ |
| Restraint stress | $P < 0.001^{**}$ | -- | $F_{4,71} = 0.834$ | $F_{4,71} = 4.046$ | $F_{4,64} = 3.353$ |
| Contact w/ female | $P < 0.001^{**}$ | $P = 0.508$ | -- | $F_{4,75} = 9.592$ | $F_{4,68} = 5.049$ |
| Contact w/male | $P < 0.001^{**}$ | $P = 0.005^{**}$ | $P < 0.001^{**}$ | -- | $F_{4,68} = 4.794$ |
| Novel object | $P = 0.008^{**}$ | $P = 0.015^*$ | $P = 0.001^{**}$ | $P = 0.002^{**}$ | -- |

Based on Shannon’s information theory (1948), we calculated the appearance probability of the cell-specific synaptic strength (Sakimoto et al, 2019). Using the appearance probability in inexperienced controls, we analyzed the appearance probability of recorded neurons one by one. Figure 6D shows cell-specific self-entropy and the visualized density distribution. The results of one-way ANOVA in individual self-entropy parameters are shown in Figure 6E (Table 20).

**Table 20.** One-way ANOVAs and *post hocs* for Figure 6E

|  | mEPSC amplitude (bit) | mIPSC amplitude (bit) |
| --- | --- | --- |
| **Overall** | $F_{4,183} = 4.057, P = 0.004^{**}$ | $F_{4,183} = 4.381, P = 0.002^{**}$ |
| Inexperienced vs Restraint | $P = 0.023^*$ | $P = 0.124$ |
| Inexperienced vs Female | $P < 0.001^{**}$ | $P = 0.013^*$ |
| Inexperienced vs Male | $P = 0.232$ | $P < 0.001^{**}$ |
| Inexperienced vs Object | $P = 0.692$ | $P = 0.105$ |
| Restraint vs Female | $P = 0.283$ | $P = 0.374$ |
| Restraint vs Male | $P = 0.943$ | $P = 0.016^*$ |
| Restraint vs Object | $P = 0.071$ | $P = 0.908$ |
| Female vs Male | $P = 0.239$ | $P = 0.116$ |
| Female vs Object | $P = 0.004^{**}$ | $P = 0.453$ |
| Male vs Object | $P = 0.075$ | $P = 0.025^*$ |
|  | mEPSC frequency (bit) | mIPSC frequency (bit) |
| **Overall** | $F_{4,183} = 2.723, P = 0.031^*$ | $F_{4,183} = 8.431, P < 0.001^{**}$ |
| Inexperienced vs Restraint | $P = 0.447$ | $P = 0.296$ |
| Inexperienced vs Female | $P = 0.226$ | $P = 0.739$ |
| Inexperienced vs Male | $P = 0.018^*$ | $P < 0.001^{**}$ |
| Inexperienced vs Object | $P = 0.459$ | $P = 0.026^*$ |
| Restraint vs Female | $P = 0.673$ | $P = 0.468$ |
| Restraint vs Male | $P = 0.118$ | $P < 0.001^{**}$ |
| Restraint vs Object | $P = 0.146$ | $P = 0.233$ |
| Female vs Male | $P = 0.239$ | $P < 0.001^{**}$ |
| Female vs Object | $P = 0.058$ | $P = 0.054$ |
| Male vs Object | $P = 0.003^{**}$ | $P < 0.011^*$ |

To examine experience specificity, we further analyzed the four self-entropy parameters in four-dimensional virtual space. We used one-way MANOVA with experience again as the between-group factor, and the results showed a significant main effect of experience (Fig. 6E; *F*_16, 551_ = 4.361, *P* < 0.001). *Post hoc* MANOVA further showed multiple differences between two specific experiences (Table 21), suggesting experience-specific information content in CA1 pyramidal neurons.

**Table 21.**
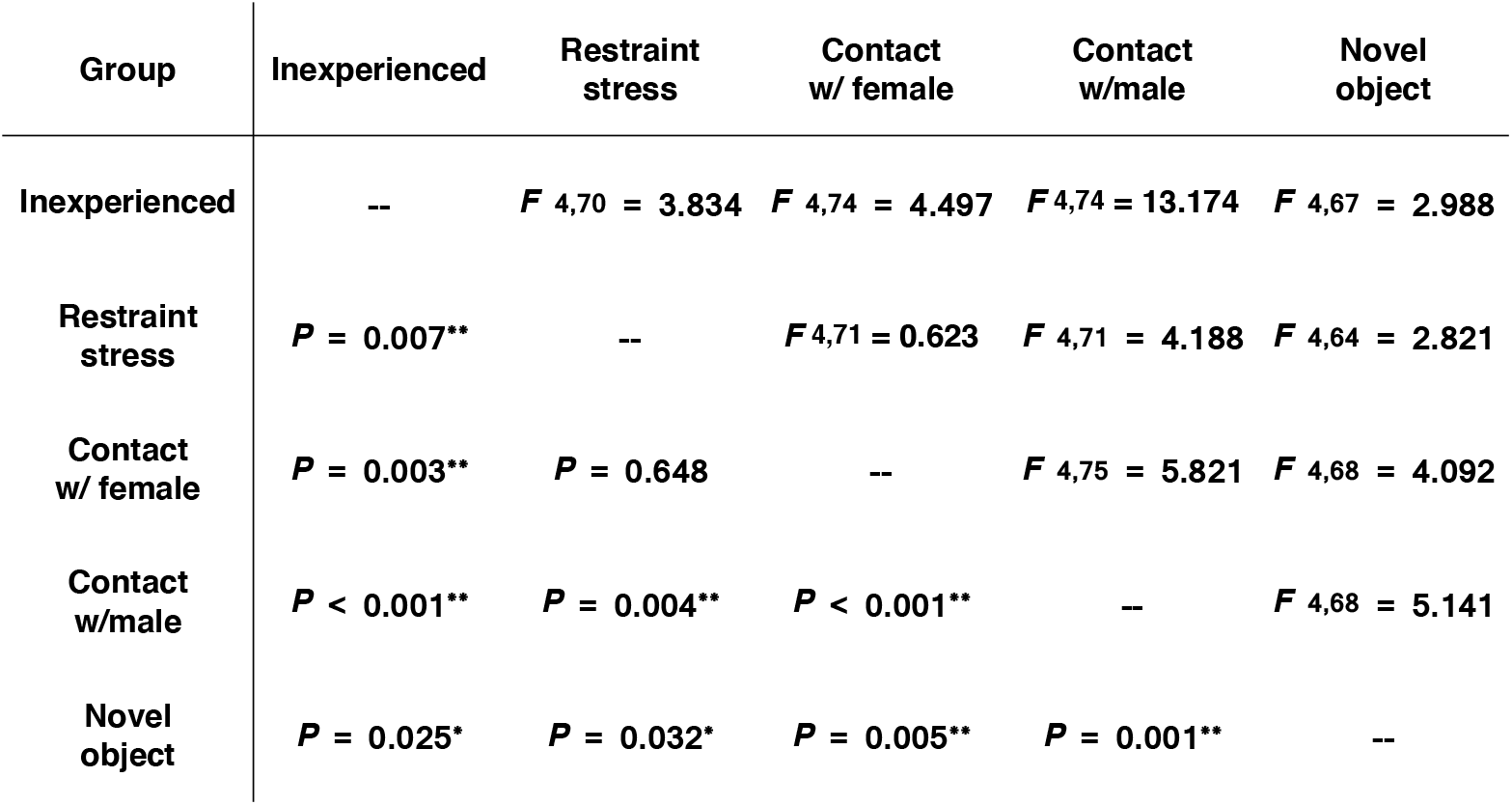
*Post hoc* MANOVAs for Figure 6E

## Discussion

Here we recorded multiple-unit firings in CA1 from freely moving male rats in a habituated home cage (Fig. 1C, Movie S1) and subjected the rats to the one of four episodic experiences for 10 min: restraint stress, social interaction with a female or male, or observation of a novel object (Fig. 1B). Based on the basal firings in the habituated home cage (Fig. 1F), we extracted three firing events of multiple-unit recording: super bursts, silent periods, and ripple-like events (Figs. 1G–I). After the onset of episodic experience, we found episode-specific intermittent generation of super bursts (Fig. 2) and frequent induction of ripple-like events with silent periods (Movie S2; Fig. 3). The four features (amplitude, duration, arc length, peaks) of these thousands of ripple-like firing events were also experience-specific (Figs. 4 and 5). Finally, *ex vivo* patch-clamp analysis showed experience-specific plasticity at excitatory/inhibitory synapses (Fig. 6).

### Synaptic plasticity

Long-term potentiation (LTP) has been considered as a synaptic model of learning and memory (Bliss & Lømo, 1972). The LTP not only enhances the presynaptic release of glutamate (Dolphin et al, 1982) but also increases the number of GluA1-containing AMPA receptors in CA1 pyramidal neurons (Shi et al, 1999, Hayashi et al, 2000). Moreover, the high-frequency stimulation at the Schaffer collaterals that induces LTP of evoked field response enhances the number of sharp-wave ripples (Buzsáki 1984, 2015). Although high-frequency electrical stimulation and concomitant pre- and postsynapse excitation can induce synaptic plasticity (Harris & Teyler, 1976; Nicoll et al, 1988), intrinsic high-frequency stimuli have not been established in animals during learning. Here we extracted spontaneous high-frequency firing events of CA1 neurons (super bursts) during and soon after the episodic stimuli.

Individual super bursts showed different firing rates and durations (Fig. 2F), and the multiple features of the super bursts were episode-specific (Table 5 and S6). The episode-specific super bursts may trigger diversification of the ripple-like events because high-frequency stimulation at CA3-CA1 synapses increases both incidence and amplitude of sharp-wave ripples in CA1 (Behrens et al, 2005). In the present study, both incident and duration of super bursts were positively correlated with the multiple features of ripple-like events (Fig. 5C), and the bilateral inactivation of basolateral amygdala by a muscimol/baclofen mixture before the restraint not only attenuated the super bursts but also blocked the diversification of ripple-like events (unpublished preliminary data). These results suggest a causal relationship between the super bursts and the ripple-like events.

### Restraint stress

Because physiological stress rapidly occludes LTP induction (Li et al, 2014, Yang et al, 2006), stress-induced synaptic plasticity has been hypothesized for decades. Regarding the morphological evidence, acute physiological stress rapidly enlarges the spine volume of CA1 pyramidal neurons within one hour (Sebastian et al, 2013), increasing spine density for more than 24 hours (Shors et al, 2004).

Restraint stress has been used as a strong stressful experience (Ulloa et al, 2010; Ciccocioppo et al, 2014; Ribeiro-Oliveir et al, 2018) that acutely induces acetylcholine release in the dorsal hippocampus (Mitsushima et al, 2008). Acetylcholine depolarizes the cell membrane, blocks subsequent hyperpolarization, and induces burst-like firings in CA1 pyramidal neurons (Cole & Nicoll, 1983). Although we previously reported that acetylcholine release triggers experience-induced synaptic plasticity in CA1 pyramidal neurons (Mitsushima et al, 2013), spontaneous highly frequent firing events had not been reported in CA1 neurons. Because restraint stress frequently induced spontaneous super bursts during the experience (Fig. 2B), these bursts may trigger or promote the rapid plasticity (< 5 min) at dorsal CA1 synapses (Sakimoto et al, 2019).

### Contact with a female

In contrast to restraint stress, contact with female may be a “positive” experience and has been used as a reward for conditioned memory (Ramirez et al, 2015; Coria-Avila, 2012). Some neurons in the dentate gyrus and basolateral nucleus of male mouse amygdala respond to a female mouse (Redondo et al, 2014; Ramirez et al, 2015). Moreover, contact with a female activates specific engram cells in the hippocampal dentate gyrus, and re-activation of the neurons reduces stress-induced depression-related behavior in male mice (Ramirez et al, 2015). Contact with a female in the present study may have been surprising for recorded males, which had never met a breedable female and exhibited long-lasting super bursts (Fig. 2E). Moreover, the diversified ripple-like events were comparable to those after the restraint (Figs. 5A and B). Although it is unclear whether this situation is applicable to humans, humans do tend to develop episodic recall of the time and location of an initial heterosexual encounter of interest (Turgenev, 1860; Carpenter & Carpenter, 1975).

### Contact with a male rat

The contact with a male was not quite as novel given that the recorded males had been reared in same sex/age group. In experiences like these, the male normally checks superiority with the unknown intruder (Whishaw & Kolb, 2005). Heterosexual contact prevents the development of conditioned same-sex partner preference in male rats, suggesting a difference between the two social memories (Ramirez-Rodriguez et al, 2017). Moreover, approximately 75% of dorsal CA1 neurons not only process their own location but also express the location of the other male rat in the same cage (Danjo et al, 2018), suggesting a large population of junction-place-cells when two male rats were housed in the same cage.

Genetically targeted inactivation of dorsal CA2 pyramidal neurons causes a pronounced loss of social memory, suggesting a critical role for CA2 neurons in socio-cognitive memory processing (Hitti et al, 2014, Stevenson et al, 2014, Caruana et al, 2012). Because CA2 pyramidal neurons have major excitatory projections to the deep layer of CA1 pyramidal neurons (Kohara et al, 2014), excitatory inputs from CA2 may contribute to forming the ripple-like firings after the contact with male. Moreover, recent optogenetic analysis in male mice also revealed a role for ventral CA1 in social memory storage (Okuyama et al, 2016). Although contextual learning failed to induce synaptic plasticity at the ventral CA1 synapses (Sakimoto et al, 2019), social experience may promote synaptic plasticity by changing the firing patterns in ventral CA1 neurons.

### Contact with a novel object

Unlike the social stimuli, the novel object did not change location. Although the induction of super bursts was unclear and inconsistent (Fig. 2B), both the object-recognition and object place–recognition tasks may increase spontaneous firings during the exploration of novel object (Munyon et al, 2014; Larkin et al, 2014). Moreover, the firing patterns in some CA1 neurons seem to be sensitive to a particular object in a particular location, and the size of place fields of dorsal CA1 neurons decreases when objects are present (Burke et al, 2011).

Regarding plastic changes, the novel object task induces a long-term decrease in the field potential response in dorsal CA1 neurons (Goh & Manahan-Vaughan, 2013). In the present study, the number of peaks in individual ripple-like events decreased in the presence of the novel object (Fig. 4H), and the postsynaptic Cl^-^ current at inhibitory synapses clearly increased (Fig. 6C). Bilateral interference of GABA_A_ receptor– mediated transmission by bicuculline or muscimol clearly blocked the encoding process in the novel object recognition task, suggesting an important role for GABA_A_ transmission in processing novel object memory at the dorsal CA1 synapses (Yousefi et al, 2013; Cohen et al, 2013).

### Features of the ripple-like event

The firing sequence during ripples replays the sequence of the location during spatial learning (Joo & Frank 2018; Foster & Wilson 2006; Diba & Buzsáki. 2007; Karlsson & Frank 2009; Davidson et al, 2009; Gupta et al, 2010; Wu et al, 2017), suggesting a role for these firings in representing the experienced sequence of the location. Moreover, spatial learning requires the represented ripples because interference of sharp-wave ripples in behaving animals impairs spatial learning (Girardeau et al, 2009; Ego-Stengel & Wilson. 2010; Jadhav et al, 2012).

Individual neurons may handle binarized firing data created by all-or-none principle (Cannon, 1922; Wiener 1961). Logarithmic relationship between the number of CA1 neurons and contextual learning performance provides the first evidence to transmit binarized data in the neurons (Mitsushima et al., 2011). Although whether past events affect ripple-like firings is unknown, operant conditioning increases the synchrony of neighboring dorsal CA1 neurons (Sakurai et al, 2013). Episode-specific changes in the ripple-like firings (Fig. 5A and B) may be a first step toward being able to identify which animal is experiencing one of the four episodes.

### Synaptic diversity in individual CA1 neurons

The question arises as to why the firing pattern changed after the experience and following episode-specific super bursts. Here we hypothesized that experience-induced plasticity at CA1 synapses may cause episode-specific changes in ripple-like firings. To evaluate plasticity at CA1 synapses, we analyzed *ex vivo* mEPSCs and mIPSCs in individual CA1 pyramidal neurons from experienced rats (Fig. 6A). Each neuron exhibited different mEPSCs and mIPSCs and frequencies at excitatory and inhibitory synapses, showing a synaptic diversity (Fig. 6B). Moreover, the distribution of the diversity differed among experiences. Although the figures are two-dimensional, experience created a specific distribution of four-dimensional plots suggesting an experience-specific synaptic diversity (Table 19). The mEPSCs and mIPSCs are thought to correspond to the response elicited by a single vesicle of glutamate or GABA (Pinheiro & Mulle, 2008), while the number of synapses affects the frequency of events. It is possible that the neuron-dependent synaptic current or number of individual neurons may contribute to creating specific firings of CA1 neurons (Fig. 7).

**Figure 7.**
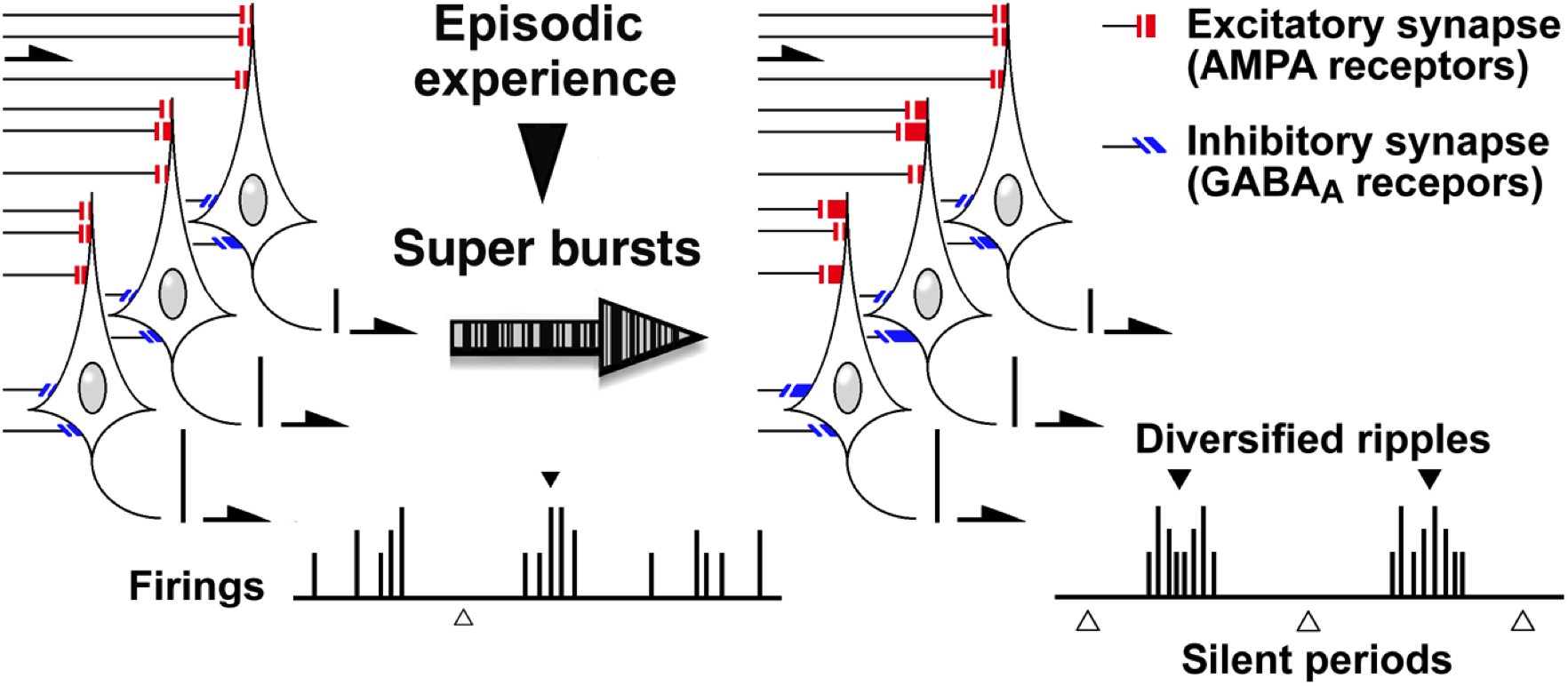
Our hypothesized early learning process in CA1 pyramidal neurons. Here we found episode-specific events of super bursts, synaptic plasticity, and ripple-like firings. Experience-induced super bursts may create specific synaptic diversity and ripple-like firings.

Both excitatory and inhibitory synaptic transmissions are important for regulating the generation of sharp-wave ripples and spiking during ripples (Buzsáki, 2015). In a slice model, sharp-wave ripples originate in CA3, propagate to CA1 and the subiculum, and require AMPA receptors. Blockade of AMPA receptors suppresses generation of sharp-wave ripples (Behrens et al, 2005; Schlingloff et al, 2014). Regarding inhibitory synapses, a single perisomatic CA3 interneuron synchronizes GABA_A_ receptor– mediated IPSCs to organize the phase-locked spiking of pyramidal cells (Ellender et al, 2010). Moreover, the form of sharp-wave ripples seems to depend on the dose of GABA_A_ receptor antagonist (Nimmrich et al, 2005; Ellender et al, 2010; Maier et al, 2003).

High frequency stimulation at CA3-CA1 synapses increases both incidence and amplitude of sharp-wave ripples in CA1 (Behrens et al, 2005), so plasticity at excitatory synapses may change the features of these ripple-like events. Especially emotional experiences (such as restraint or contact with a female) were associated with clear spontaneous super bursts (Fig. 2B), strengthened excitatory synapses (Fig. 6C), and increased incidence (Fig. 3F), amplitude (Fig. 4B), and duration of ripple-like events (Fig. 4D). The longer duration may result from strengthened excitatory synapses because the mixture of AMPA/NMDA receptor antagonists decreases the duration of evoked ripple-like events (Schlingloff et al, 2014). Recent optogenetic approaches in CA1 pyramidal neurons have shown that artificially prolonging sharp-wave ripples improves working memory performance, whereas aborting the late part of ripples decreases it (Fernández-Ruiz et al, 2019). These reports together with the present results suggest a definitive role for ripple-like events in processing the experiences.

### Conclusion

Figure 7 shows our hypothesized early learning process in CA1 pyramidal neurons. Experience induces episode-specific super bursts that may promote episode-specific synaptic diversity and ripple-like events. If this conceptualization is correct, it may be possible to decipher encrypted experience through the deep learning of the orchestrated ripple-like firings and synaptic plasticity in multiple CA1 neurons. In addition, understanding the orchestrated ripple-like firings and plasticity in multiple CA1 neurons may help specify signal failures in multiple cognitive disorders.

## Author Contributions

JI and DM performed the experiments. TT, JI, and DM analyzed firing events. DM and JI wrote the manuscript. DM organized the study, and all authors reviewed the manuscript.

## Acknowledgments

The authors thank Ryo Sato for the analysis of ripple-like events. This project is supported by the Grant-in-Aid for Scientific Research B (Grants 16H05129 and 19H03402 to DM), Scientific Research C (Grants 26350988 and 17K01987 to JI, Grant 25460314 to DM), and Scientific Research in Innovative Areas (Grant 26115518 to DM) from the Ministry of Education, Culture, Sports, Science and Technology of Japan. The authors declare no conflicts of interest.

